# GWAS and multi-omics uncover the *SLC6A3* locus for canine stranger-directed social anxiety and implicate *NRIP3* in dopaminergic network regulation

**DOI:** 10.64898/2026.08.10.744065

**Authors:** Huairen Zhao, Judong Li, Jing Li, Sinan Wang, Ye Liu, Yaoyue Wu, Fakun Zhang, Shurun Zhang, Kang Huang, Shihua Xue, Jiusheng Wan, Yun Yu, Ya-Ping Zhang

## Abstract

The social anxiety disorder (SAD) is one of the most common mental health disorders, often developing in adolescence. It can severely impair educational achievement, career progression, and social functioning. Certain individuals within the Chinese Kunming dog (CKD) population display spontaneous, SAD-like behaviors towards strangers, offering a valuable model for genetic investigation. Using whole-genome sequencing data, we conducted a genome-wide association study on this SAD-like behavior in 100 CKDs, identifying several genes previously linked to human psychiatric disorders. Notably, *SLC6A3* reached genome-wide significance, whereas *CADPS2* exceeded only the suggestive threshold and is therefore considered a network-supported candidate locus. By integrating weighted gene co-expression network analysis with public RNA-seq data from the VTA of 38 mice (after quality control), we further showed that *Slc6a3* and *Cadps2* are part of a co-expression network module associated with the dopamine system. These findings suggest that this specific module may play a functional role in the stranger-directed anxiety-related behaviours in CKDs. Furthermore, we identified *Nrip3* as a key candidate transcriptional coregulator in this specific module. Subsequent virus-mediated overexpression and behavioural experiments showed that *Nrip3* regulates gene expression within this network and induces a spectrum of behavioural alterations in mice, including social avoidance, anhedonia, and increased locomotor activity. Collectively, this study highlights the involvement of dopamine system genes in canine social anxiety and suggests *Nrip3* as an upstream regulator of *Slc6a3* associated with anxiety. The identification of conserved anxiety-related signatures between humans and dogs further supports the dog as a valuable translational model for psychiatric research.

## Introduction

Social anxiety disorder (SAD) is a common and debilitating psychiatric disorder with an estimated lifetime prevalence rate of 12.1% (Grison et al., 2005). It is characterized by a marked and persistent fear of one or more social situations (e.g., talking to a stranger or peer) or performance activities (e.g., giving a speech) in which the person is exposed to unfamiliar people, or where they may face possible scrutiny by others (Bandyopadhyay Prasanta et al., 2014). Although prior genome-wide association study (GWAS) analyses have yielded a series of loci and genes associated with SAD (Baba et al., 2022; Stein et al., 2017), the heterogeneity and polygenic architecture of the disorder have posed substantial challenges to comprehensively unravelling its full genetic architecture.

Dogs have emerged as a compelling and underutilized animal model for dissecting the genetics of human psychiatric traits, including anxiety-related disorders (Morrill et al., 2023; Zapata et al., 2020). Accumulating evidence suggests that dogs and humans have undergone convergent evolution in social cognition and behavior, shaped by millennia of close cohabitation and cooperative interactions (Miklósi & Topál, 2013; Range & Virányi, 2015). Previous studies revealed that dogs also exhibit fearful or anxious behavior, and genetic investigations suggest a similar genetic background between canine anxiety and some human psychiatric diagnoses characterized by anxiety and fear (Overall, 2000; Sarviaho et al., 2019; Yu et al., 2026). The unique population history of domestic dogs, shaped by intense artificial selection during breed formation, has resulted in reduced genetic diversity within breeds and extended linkage disequilibrium (LD) blocks. This LD structure, combined with homogeneous breed backgrounds, facilitates genetic mapping with relatively small sample sizes compared to human populations (Halo et al., 2021; Lindblad-Toh et al., 2005; Meadows et al., 2023; Sutter et al., 2004; Wang et al., 2021). This population history has also resulted in a high frequency of genetic disease in some dog breeds (Donner et al., 2023; Dutrow et al., 2022), including anxiety and fear (Overall, 2000; Sarviaho et al., 2019). Despite these compelling reasons to use dogs for scientific discovery relevant to both canine and human biology, dogs have been underused as a model organism to date (Wallis et al., 2025).

Inferring causal genes from GWAS findings alone in dogs is challenging and typically requires additional functional validation through *in vitro* and *in vivo* experiments. A powerful complementary approach involves integrating GWAS results with multi-omics data, including transcriptomic data (Kreitmaier et al., 2023; Wang et al., 2018; Xu et al., 2025). Bulk and single-cell transcriptomic datasets from mice, which are massively produced and available. Given the high evolutionary conservation of gene functions across mammals, across-species omics integration enables reliable inference of gene function and predicted regulatory relationships (Reaume & Sokolowski, 2011). Specifically, combining GWAS-derived candidate genes with gene co-expression networks allows precise prioritization of disease-associated genes: genes exhibiting coordinated expression patterns frequently participate in shared biological pathways, respond to common regulatory signals, or are co-regulated by overlapping transcription factor modules (Paci et al., 2021). Furthermore, by simultaneously inferring upstream regulators and downstream effectors from transcriptomic analysis (Van de Sande et al., 2020), this approach extends the scope of candidate gene discovery well beyond the limitations of initial GWAS signals, providing an extended view of the disease architecture.

The Chinese Kunming Dog (CKD), developed in China since the 1950s by hybridizing German Shepherds with indigenous dogs from Kunming, has been recognized as a new breed by the Chinese National Livestock and Poultry Genetics Commission since 2007 (Huang et al., 2024). Preliminary behavioral assessments indicate that a subset of CKDs exhibits intense, spontaneous fear and anxiety when encountering unfamiliar individuals. Manifestations of this temperament include avoidant behavior and, in some instances, pronounced tremors when approached by strangers. In this study, we collected 18 CKDs with typical social-anxiety behavior and 82 normal individuals, and obtained whole-genome sequencing (WGS) data of them. We then performed a GWAS on this cohort of 100 CKDs and identified novel loci on chromosome 14, 27, and 34. Integrating spatial-temporal and bulk transcriptomic data from mice, we showed that (a genome-wide significant locus) *SLC6A3* and *CADPS2* (a suggestive, network-supported candidate locus) are located in the same co-expression module and are both implicated in dopaminergic synaptic function. We also identified a candidate upstream modulator gene of this gene module, *Nrip3*. Subsequently, via viral overexpression of the *Nrip3* in the ventral tegmental area (VTA) of mice and behavioral tests, we showed that this *Nrip3* overexpression is associated with alterations in behavior, including social behavior, spontaneous locomotor behavior, and anhedonia. Bulk transcriptomic analysis verified that this gene indeed regulates expression of genes including *Slc6a3* and *Cadps2*, and other genes involving in the co-expression module.

## Results

### Genetic dissection of social anxiety-like behavior in Chinese Kunming dogs

To elucidate the genetic architecture of SAD-like behavior in CKDs, we generated 20× coverage WGS data from peripheral blood samples. Following quality control to filter for high-quality individuals and variants, the study included 100 CKDs (comprising a case group of 18 individuals exhibiting SAD-like behavior and a control group of 82 behaviorally normal individuals) (Supplementary Table S1), yielding 5,624,028 single-nucleotide polymorphisms (SNPs) and insertions/deletions (InDels). To control for population stratification and cryptic relatedness, we performed principal component analysis (PCA) using 174,290 LD-pruned (r²<0.2) variants and included the first two principal components (PC1, explaining 19.5% of the variance; PC2, explaining 12.7%; Supplementary Figure S1) together with sex as covariates in a linear mixed model (LMM) implemented in GEMMA (Zhou & Stephens, 2012). Additionally, A VanRaden standardized genomic relatedness matrix (GRM) was used to account for sample relatedness. The quantile-quantile (QQ) plot showed no systematic deviation from the null expectation (Supplementary Figure S2), with a genomic inflation factor of λGC = 1.07, indicating adequate control of population stratification. Principal component analysis confirmed that cases and controls were well-mixed across the first two axes of genetic variation.

The GWAS identified two genomic peaks exceeding the Bonferroni genome-wide significance threshold, located on chromosome 34 within the *SLC6A3* gene and on chromosome 27, respectively. An additional peak on chromosome 14 exceeded the suggestive threshold but did not reach genome-wide significance (Figure 1A). To assess the robustness of these associations given the modest sample size, we performed three sensitivity analyses (Supplementary Tables S2–S4). First, a leave-one-case-out analysis was conducted by iteratively removing each of the 18 case individuals and re-running the full GEMMA LMM (n = 99 per iteration). Across all 18 iterations, the effect direction (β) at each lead SNP remained consistently negative, and the corresponding *p* remained below the suggestive threshold of 1 × 10⁻⁵ (Supplementary Table S2). Second, permutation testing (100,000 phenotype permutations, PLINK --mperm) yielded an empirical *p* (EMP2) of 0.013 for the chromosome 34 lead SNP, providing additional empirical support for this locus. The chromosome 14 (EMP2 = 0.581) and chromosome 27 (EMP2 = 0.282) loci did not reach permutation significance under the simpler allelic χ² test, indicating stronger permutation support for the Chr34/*SLC6A3* signal and more limited permutation support for the other two loci (Supplementary Table S3). Third, the alternate (risk) allele frequencies were 1.4- to 5.1-fold higher in cases than in controls (Chr14: 83.3% vs. 37.8%; Chr27: 52.8% vs. 13.4%; Chr34: 55.6% vs. 11.0%;), and all 95% confidence intervals excluded zero. Because GEMMA -lmm models binary phenotypes using a linear mixed model, we report β and its 95% CI rather than odds ratios. Effect sizes (β), standard errors, 95% confidence intervals, and allele frequencies in cases versus controls were estimated for all three lead SNPs (Supplementary Table S4). Given the small sample size (*n* = 100), we present these findings as candidate signals warranting replication in independent cohorts. Together, the genome-wide significant signal at Chr34/*SLC6A3*, stable effect directions across leave-one-out iterations, permutation support for the chromosome 34 locus, and biologically plausible candidate genes support prioritization of these loci as candidate signals requiring independent replication.

**Figure 1.**
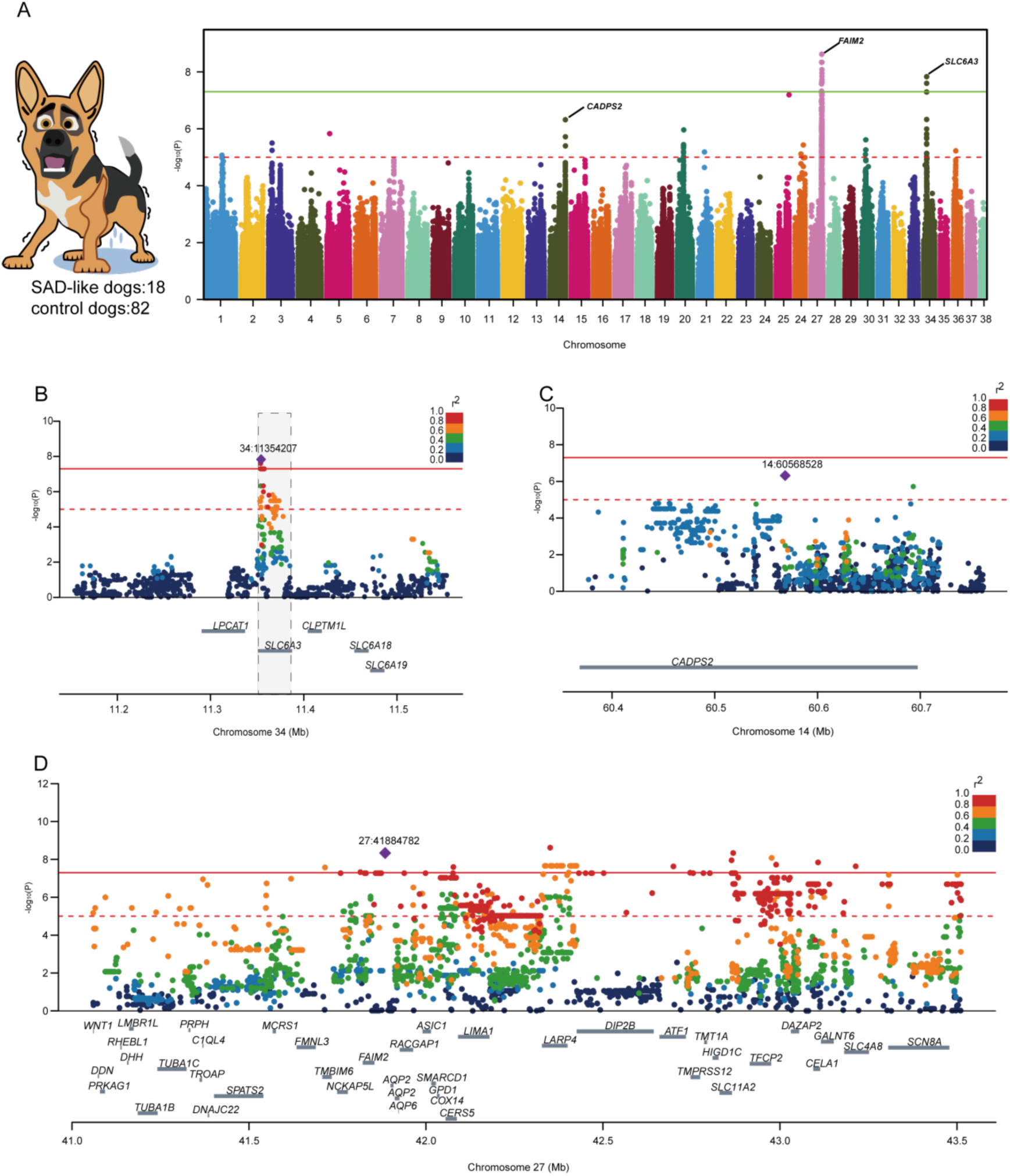
GWAS of social anxiety-like behavior in Chinese Kunming dogs. (A) Manhattan plot showing the results of the genome-wide association study for SAD-like behavior in 100 Chinese Kunming dogs (18 cases, 82 controls). (B) Regional association plot for the chromosome 34 locus. The defined ROI (comprising SNPs in high LD, r² ≥ 0.7, with the lead SNP) maps entirely within the *SLC6A3* gene, which reaches genome-wide significance. (C) Regional association plot for the chromosome 14 locus. Two variants surpass the suggestive significance threshold, identifying *CADPS2* as a suggestive, functional supported candidate gene in this region. (D) Regional association plot for about 2.5 Mb ROI on chromosome 27. The extensive LD block encompasses 41 known protein-coding genes.

For each significant signal, we defined the region of interest (ROI) as the continuous span of SNPs in high linkage disequilibrium (LD, r² ≥ 0.7) with the lead SNP (Alex et al., 2025). Notably, the ROI located on chromosome 34 is entirely within the *SLC6A3* gene (Figure 1B). *SLC6A3* encodes the dopamine transporter (DAT), which primarily reuptakes dopamine from the synaptic cleft into dopaminergic neurons, thereby modulating the intensity and temporal dynamics of dopamine signaling (M. E. Reith et al., 2022). Previous studies have shown that genetic loss or targeted knockdown of *Slc6a3* in mice results in significantly reduced anxiety behaviors (Bahi & Dreyer, 2019; Petri et al., 2024). Furthermore, DAT serves as a crucial pharmacological target for treating various psychiatric and neurological disorders, including depression, attention-deficit/hyperactivity disorder (ADHD), and Parkinson’s disease (Nepal et al., 2023; M. E. Reith et al., 2022).

The peak on chromosome 14 (60.4–60.7 Mb) that exceeds the suggestive threshold but does not reach genome-wide significance (Figure 1C). Variants surpassing this suggestive threshold are located within the *CADPS2* gene, which we designate a functional supported candidate rather than a genome-wide significant locus. It encodes a calcium-dependent regulator of exocytosis that complements DAT (dopamine transporter) at dopaminergic synapses (Iguchi, Katsuzawa, Saruta, Sadakata, Kobayashi, Sato, Sato, Sano, Maezawa, Shinoda, et al., 2024). Consistent with this synaptic and neurosecretory role, *Cadps2* mutant mouse models have been reported to show impaired social interaction and anxiety-related behavioral alterations in unfamiliar environments, and oxytocin-neuron-specific *Caps2* conditional knockout mice exhibit impaired social interaction and recognition behavior (Fujima et al., 2021; Sadakata et al., 2012; T. Sadakata et al., 2007). Both the *SLC6A3* and *CADPS2* genes are highly expressed in the substantia nigra pars compacta (SNc) and Ventral Tegmental Area (VTA) according to public spatial single-cell transcriptomic data from the mouse brain (Supplementary Figure S3, S4).

Notably, chromosome 27 harbors a large ROI spanning about 2.5 Mb, which contains 41 protein-coding genes (Figure 1D). The lead variant in this region is located within the intergenic region 30 kb downstream of *FAIM2* (Fas apoptotic inhibitory molecule 2). *FAIM2* is a lysosome-localized member of the transmembrane BCL2-associated X protein (BAX) inhibitor motif-containing (TMBIM) family and plays critical roles in nervous system development, anti-apoptosis, and autophagy regulation (Hong et al., 2020). In addition, *ASIC1* (acid-sensing ion channel 1), also located in this locus, has been implicated in emotional regulation (Shi et al., 2023). Furthermore, *WNT1*, encoding a key regulator of midbrain dopaminergic neuron development (Prakash et al., 2006), and *SCN8A*, a high-confidence autism spectrum disorder gene (SFARI Score 1) associated with epilepsy and ADHD (Johannesen et al., 2022), are also located in this region. Among the 41 protein-coding genes in this region, *AQP6* and *CERS5* are also known to perform important functions in the central nervous system (Han et al., 2016; Zhang et al., 2025). Given the function of these genes, it is difficult to infer the candidate causal in this genomic region. We therefore opt to prioritize regions with more discrete genetic architectures for immediate validation, deferring a detailed analysis of this region to future investigations.

### *Slc6a3* and *Cadps2* triggers highly similar downstream transcriptional remodeling and pathway enrichments

Given that both *Slc6a3* and *Cadps2* are highly specific to dopaminergic neurons (Bonora et al., 2014; Iguchi, Katsuzawa, Saruta, Sadakata, Kobayashi, Sato, Sato, Sano, Maezawa, & Shinoda, 2024), we selectively isolated this cell population for subsequent *in silico* knockout analyses to explore the predicted functions of these two genes (Figure 2A). Using canonical cell-type-specific marker genes (dopaminergic neurons: *Slc6a3, Th, Drd2,* and *Nr4a2; Slc6a3* shown as the representative marker in Figure 2B; Supplementary Figure S5), we identified eight major cell populations, including glutamatergic neurons, microglia, dopaminergic neurons, GABAergic neurons, NG2-glia/oligodendrocyte precursor cells (OPCs), oligodendrocytes, astrocytes, and Schwann cells (Figure 2C).

**Figure 2.**
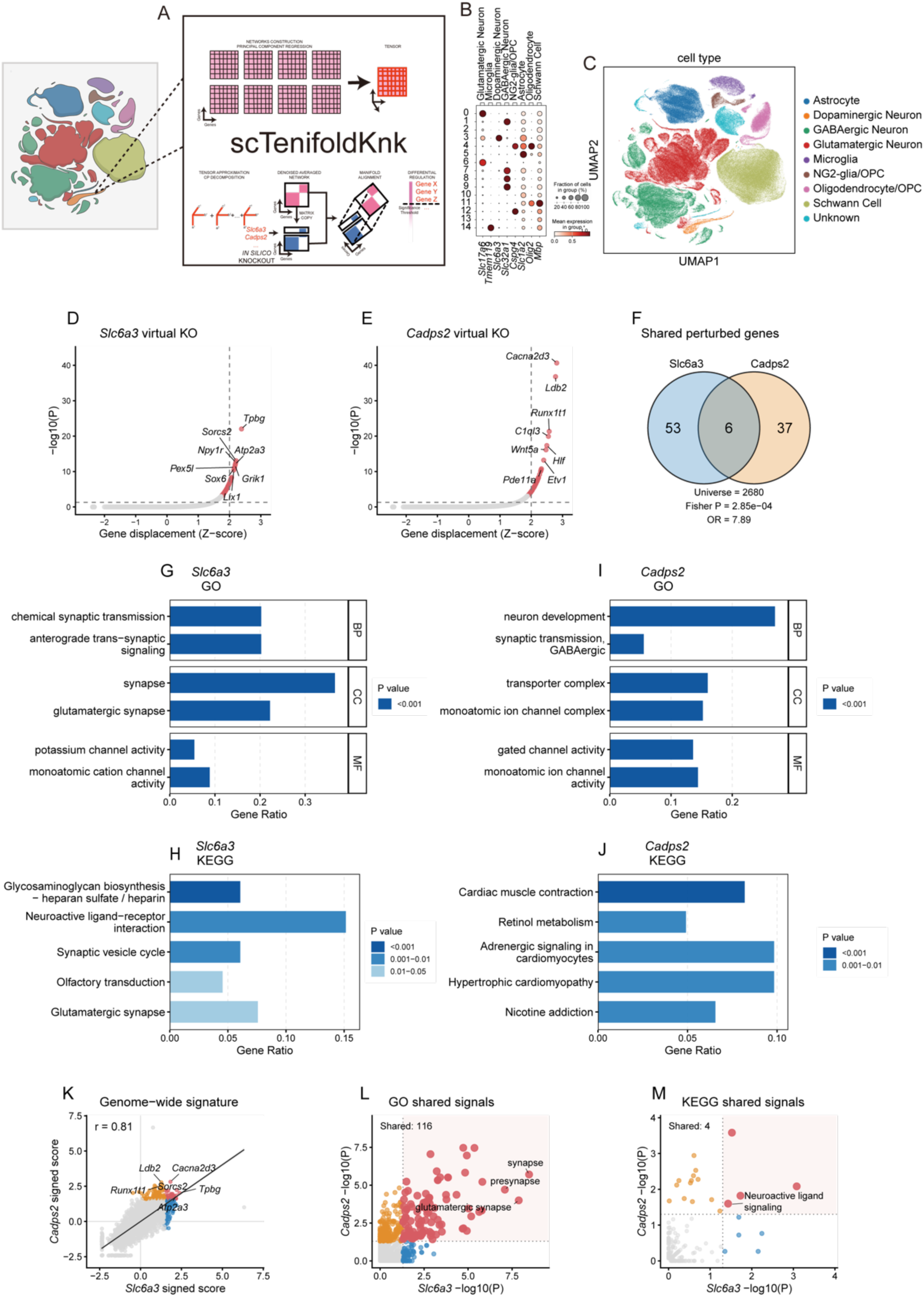
Single-cell transcriptomic profiling and *in silico* knockout analyses reveal convergent predicted network perturbations for *Slc6a3* and *Cadps2* in dopaminergic neurons. (A–C) Identification and isolation of VTA cell populations. (A) Schematic of the analytical workflow for isolating specific cell populations for subsequent *in silico* knockout analysis. (B) Expression dot plot of canonical cell-type-specific marker genes used for cluster annotation. (C) Dimensionality reduction plot revealing eight major cell populations: astrocytes, dopaminergic neurons, GABAergic neurons, glutamatergic neurons, microglia, NG2-glia/OPCs, oligodendrocytes/OPC, and Schwann cells. (D) *in silico* knockout of *Slc6a3* in the dopaminergic neuron subpopulation significantly alters of multiple genes, notably *Tpbg*, *Sorcs2*, *Npy1r*, *Atp2a3*, *Grik1*, and *Sox6*. (E) *in silico* knockout of *Cadps2* showing robust transcriptomic fluctuations in downstream genes, including *Cacna2d3*, *Ldb2*, *Runx1t1*, and *C1ql3*. (F) Overlap analysis showing notable concordance between the predicted perturbed gene alterations induced by the *in silico* knockouts of *Slc6a3* and *Cadps2,* statistical background (universe) defined as the shared set of 2,680 genes evaluated in both *Slc6a3* and *Cadps2* scTenifoldKnk *in silico* knockout outputs. (G) Gene ontology (GO) enrichment of significantly perturbed genes (*p* < 0.05) following *Slc6a3 in silico* knockout, categorized by biological process (BP), cellular component (CC), and molecular function (MF). (H) GO enrichment analysis following *Cadps2 in silico* knockout (*p* < 0.05). (I) KEGG pathway analysis for *Slc6a3 in silico* knockout targets (*p* < 0.05). (J) KEGG pathway analysis for *Cadps2 in silico* knockout. (K) Scatter plot comparing the full gene perturbation signatures (signed Z-scores) between *Slc6a3* and *Cadps2 in silico* knockouts across all 2,680 commonly tested genes, revealing a high correlation in the direction and magnitude of gene-level perturbation (Pearson r = 0.81, Spearman r = 0.82). (L) Scatter plot comparing GO enrichment significance (-log10 *p*) between *Slc6a3* and *Cadps2 in silico* knockouts; red points indicate GO terms significant (*p* < 0.05) in both knockouts. (M) Scatter plot comparing KEGG pathway enrichment significance (-log10 *p*) between *Slc6a3* and *Cadps2 in silico* knockouts; red points indicate KEGG pathways significant (*p* < 0.05) in both knockouts.

Within the defined dopaminergic neuron subpopulation, we employed the single-cell network inference algorithm scTenifoldKnk to simulate an *in silico* knockout of the candidate genes *Slc6a3* and *Cadps2*. The results showed that the *in silico* knockout of *Slc6a3* significantly altered the expression of multiple downstream genes, prominently including *Tpbg*, *Sorcs2*, *Npy1r*, *Atp2a3*, *Grik1*, and *Sox6* (Figure 2D). Similarly, the *in silico* knockout of *Cadps2* with the most robustly fluctuating genes, including *Cacna2d3*, *Ldb2*, *Runx1t1*, and *C1ql3* (Figure 2E). The predicted perturbed genes alterations induced by *in silico* knockout of these two genes exhibited notable concordance and statistically significant overlap (Fisher’s exact test, *p* = 2.85 × 10⁻⁴, OR = 7.89; universe = 2,680 genes shared between the *Slc6a3* and *Cadps2* scTenifoldKnk differential regulation outputs; *Slc6a3*: 59 significant genes at *p*.adj < 0.05, *Cadps2*: 43 significant genes at *p*.adj < 0.05, overlap: 6 genes — *Tpbg, Sorcs2, L3mbtl4, Zfp804b, Unc5d, and Cacna2d3*) (Figure 2F).

We extracted the genes significantly perturbed after *in silico* perturbation of *Slc6a3*, respectively, and performed gene ontology (GO) enrichment analysis using the shared set of 2,680 genes evaluated in both scTenifoldKnk outputs as the statistical background (universe) (Figure 2D & 2G). The results indicated that 182 GO terms reached significance (*p* < 0.05), including 94 biological process (BP), 45 cellular component (CC), and 43 molecular function (MF) terms. In the cellular component (CC) category, the most significantly enriched terms included synapse (*p* = 3.41 × 10⁻⁹), glutamatergic synapse (*p* = 1.32 × 10⁻⁸), presynapse (*p* = 8.13 × 10⁻⁸), synaptic membrane (*p* = 1.55 × 10⁻⁶), and monoatomic ion channel complex (*p* = 4.35 × 10⁻⁶). In the biological process (BP) category, chemical synaptic transmission (*p* = 1.67 × 10⁻⁶), anterograde trans-synaptic signaling (*p* = 1.67 × 10⁻⁶), and nervous system process (*p* = 9.41 × 10⁻⁶) were significantly enriched. In the MF category, enriched terms included potassium channel activity (p = 3.23 × 10⁻⁴), monoatomic cation channel activity (p = 3.87 × 10⁻⁴), heparan sulfate sulfotransferase activity (p = 4.25 × 10⁻⁴), channel activity (p = 5.27 × 10⁻⁴), and monoatomic ion channel activity (p = 6.04 × 10⁻⁴). KEGG pathway analysis showed enrichment (*p* < 0.05) in neuroactive ligand-receptor interaction (*p* = 5.71 × 10⁻³), synaptic vesicle cycle (*p* = 7.01 × 10⁻³), glutamatergic synapse (*p* = 2.05 × 10⁻²), and neuroactive ligand signaling (*p* = 3.71 × 10⁻²) (Figure 2H).

We performed enrichment analysis on the *Cadps2 in silico* knockout and its responsive differentially expressed genes using the same shared 2,680-gene universe (Figure 2E & 2I). GO results yielded 376 terms at nominal significance (*p* < 0.05). In the cellular component (CC) category, transporter complex (*p* = 3.40 × 10⁻⁸), monoatomic ion channel complex (*p* = 3.45 × 10⁻⁸), synapse (*p* = 1.96 × 10⁻⁶), and synaptic membrane (*p* = 6.06 × 10⁻⁶) were significantly enriched. In the molecular function (MF) category, gated channel activity (*p* = 8.92 × 10⁻⁸), monoatomic ion channel activity (*p* = 1.20 × 10⁻⁶), and calcium ion transmembrane transporter activity (*p* = 1.52 × 10⁻⁵) were prominently enriched, pointing to intracellular calcium dyshomeostasis and disruption of calcium-dependent exocytosis. KEGG pathway analysis showed enrichment (*p* < 0.05) in cardiac muscle contraction (*p* = 2.62 × 10⁻⁴), pathways of neurodegeneration – multiple diseases (*p* = 7.61 × 10⁻³), Alzheimer disease (*p* = 1.81 × 10⁻²), calcium signaling pathway (*p*= 4.04 × 10⁻²), and neuroactive ligand signaling (*p* = 2.52 × 10⁻²) (Figure 2J).

Additionally, we compared the full gene perturbation signatures between *Slc6a3* and *Cadps2 in silico* perturbation outputs across the shared set of 2,680 evaluated genes. The gene-level perturbation directions were highly correlated (Pearson r = 0.81, Spearman r = 0.82; Figure 2K), indicating that both *in silico* knockouts converge on similar downstream gene programs. However, at the pathway level, shared enrichment signals between the two *in silico* knockouts were more limited: while several GO terms related to synaptic transmission were co-enriched (Figure 2L), only a few KEGG pathways (e.g., glycosaminoglycan biosynthesis, cardiac muscle contraction, neuroactive ligand signaling) reached significance (Figure 2M).

### WGCNA identifies *Slc6a3* and *Cadps2* within the same co-expression network module in the mouse VTA

To further explore the potential synergistic regulatory mechanisms between *Slc6a3* and *Cadps2*, we performed weighted gene co-expression network analysis (WGCNA) on bulk transcriptome data from public mouse VTA samples (*n* = 38 after quality control). Module assignment and module eigengene clustering revealed that *Slc6a3* and *Cadps2* were co-clustered into the same co-expression module (the green module) (Figure 3A). Linear regression analysis further showed a highly significant positive correlation between the expression levels of these two genes (Pearson r = 0.96, Spearman ρ = 0.94; *p* < 0.001) (Figure 3B). To assess whether this co-expression reflects genuine gene-level co-regulation rather than shared variation in dopaminergic neuron proportions, we additionally computed the *Slc6a3–Cadps2* correlation within the annotated dopaminergic neuron subpopulation (*n* = 3,353 cells), using the raw gene-level count matrix from the 10Xv3 single-cell RNA-seq dataset before normalization or log transformation, with cell type held constant by subsetting to annotated dopaminergic neurons. The correlation remained significant (Pearson r = 0.65, Spearman ρ = 0.76; both *p* < 10⁻²⁰⁰; Supplementary Table S6; Supplementary Figure S6), confirming that the co-expression is not merely a byproduct of dopaminergic neuron abundance variation, although the reduced magnitude relative to bulk (r = 0.96) indicates that tissue-level co-expression partly reflects cell-type composition.

**Figure 3.**
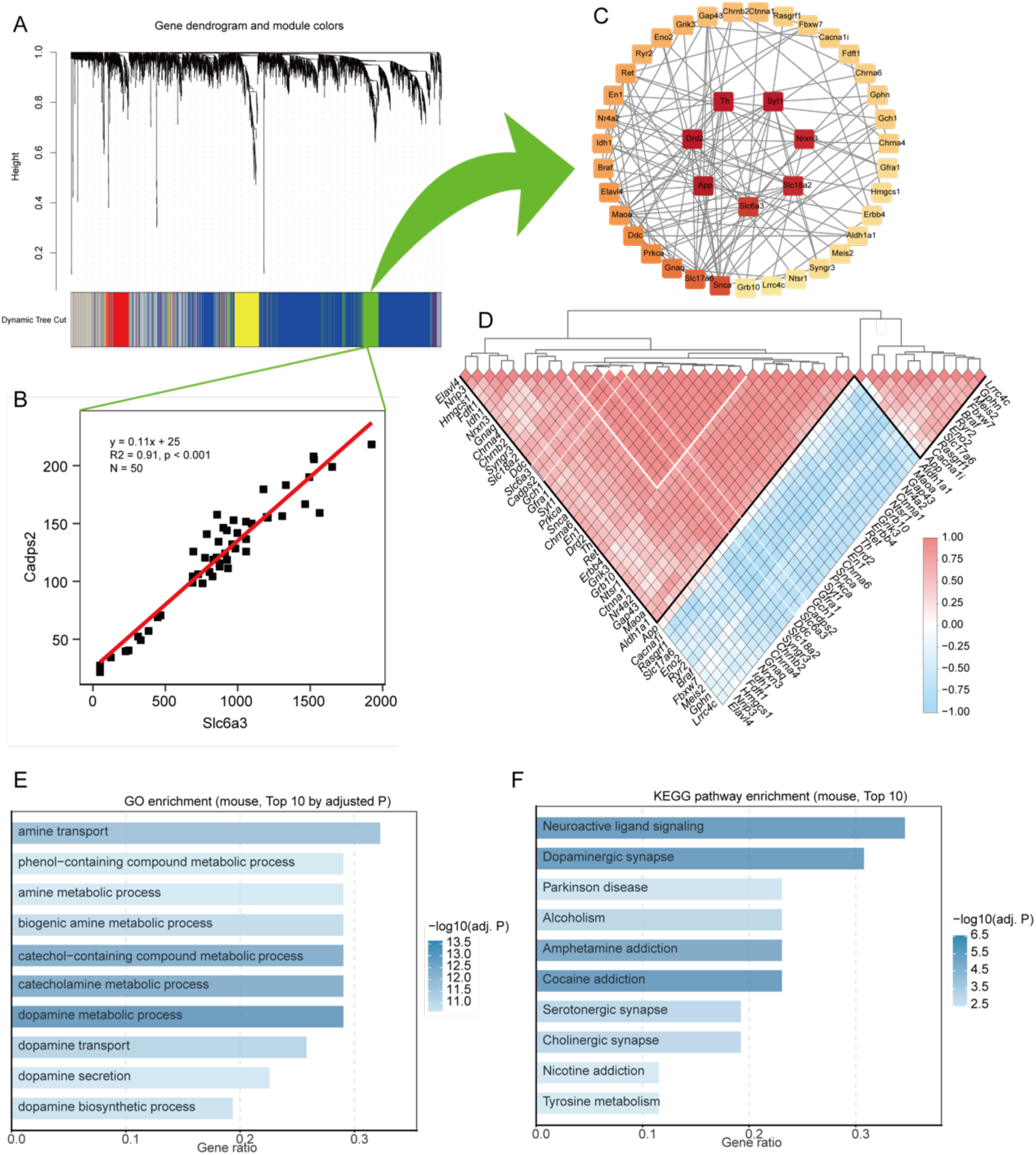
Co-expression and synergistic regulatory network of *Slc6a3* and *Cadps2* in the ventral tegmental area (VTA). (A) Weighted gene co-expression network analysis (WGCNA) of bulk RNA-seq data from mouse VTA samples (*n* = 50). Module assignment and eigengene clustering reveal the co-localization of *Slc6a3* and *Cadps2* within the green co-expression module. (B) Linear regression analysis demonstrates a highly significant positive correlation between the expression profiles of *Slc6a3* and *Cadps2* (*p* < 0.001, R² = 0.91). (C) Protein-protein interaction (PPI) network constructed from genes within the green module. Topological analysis identifies 43 principal genes, including 7 critical hub genes with high connectivity scores: *App*, *Drd2*, *Th*, *Syt1*, *Nrxn3*, *Slc18a2*, and *Slc6a3*. (D) Correlation matrix of the 43 principal genes within the PPI network. A highly correlated cluster encompasses *Slc6a3*, *Cadps2*, and dopamine-system-associated genes (*Drd2*, *Slc18a2*, *Ddc*), with 31 genes exhibiting positive correlations within the network. (E) Gene Ontology (GO) and (F) KEGG pathway enrichment analyses of the dopamine metabolism, catecholamine metabolism, amine transport, neuroactive ligand signaling, and dopaminergic synapse-related functions.

To further explore the functional nodes within this co-expression module, we extracted the green module gene set and constructed a protein-protein interaction (PPI) network analyze (Figure 3C). According to topological analysis and centrality evaluation of the network, we identified 43 principal genes, including 7 critical hub genes, *App, Drd2, Th, Syt1, Nrxn3, Slc18a2,* and *Slc6a3,* exhibiting exceptionally high connectivity scores. These 43 genes exhibit a high degree of co-expression within this module, with 31 genes showing strong positive correlations among each other (Figure 3D). Notably, 6 of the 7 critical hub genes also exhibited strong positive correlations with one another, and this highly coordinated expression pattern was consistently observed at the single-cell level (Supplementary Figure S6). Simultaneously, *Slc6a3* and *Cadps2* were identified within the same co-expression module, which also encompasses *Drd2*, *Slc18a2*, and *Ddc* genes that are fundamentally linked to the dopaminergic system. Subsequently, GO and KEGG pathway enrichment analyses were performed on the 31 positive correlation node genes of this WGCNA/PPI network (Figure 3E & 3F). The enrichment background was defined from the same mouse VTA expression dataset used for WGCNA (GSE228031), after gene-symbol processing, removal of duplicated symbols, and low-expression filtering using the WGCNA preprocessing threshold rowMeans > 1. This yielded 13,726 expressed background genes, of which 13,566 were successfully mapped to mouse Entrez IDs and used as the effective GO/KEGG universe. Using this WGCNA-filtered background, the candidate genes remained enriched for dopamine metabolism, catecholamine metabolism, amine transport, neuroactive ligand signaling, and dopaminergic synapse-related pathways. To ascertain whether the synergistic expression pattern of *Slc6a3* and *Cadps2* is evolutionarily conserved across species, we integrated a public transcriptomic dataset comprising 24 human VTA samples for WGCNA. The results demonstrated that these two genes were assigned to the same co-expression network (the yellow module) in the human VTA, with identical and robust assignments (Supplementary Figure S7). Thus, the observed synergistic expression relationship between *Slc6a3* and *Cadps2* appears to be conserved between mice and humans. Despite limited canine transcriptomic data that preclude direct validation, cross-species conservation supports the biological plausibility of similar synergistic roles of these genes in the dog brain, although direct canine transcriptomic validation is still needed.

### *Nrip3* orchestrates the expression of genes involved the co-expression network module in the VTA

It is well established that both *Slc6a3* and *Cadps2* genes play crucial roles in dopaminergic synapse-related functions. Leveraging conserved midbrain dopamine system (Demin et al., 2022; Fiorenzano et al., 2024; Kamath et al., 2022b; Suresh et al., 2023), we used mouse spatial single cell transcriptomic data to further explore expression correlation between *Slc6a3* and *Cadps2*. Initially, we applied the gsMap (Song et al., 2025) to improve the spatial resolution of the mouse whole-brain spatial transcriptomic data. Subsequently, by calculating the Pearson correlation between the gene spatial scores (GSSs) of all genes across the genome and the GSS of *Slc6a3* specifically within the SNc and VTA, we constructed a *Slc6a3*-anchored expression correlation landscape (Figure 4A; Supplementary Table S6). The results showed a highly significant positive correlation between the spatial expression patterns of *Cadps2* and *Slc6a3* (Pearson r = 0.50, Spearman ρ = 0.53), where these two genes are not only co-enriched within the SNc/VTA (Supplementary Figure S3) but also show strict synchronous co-expression across the cell populations of these regions. Furthermore, we observed that *Nrip3* gene ranked third in the expression correlation table (Pearson r = 0.67, Spearman ρ = 0.71 with Slc6a3; Supplementary table S6), which is also in the co-expression network module identified by the WGCNA. This gene encodes the nuclear receptor interacting protein 3 (NRIP3), acting as a transcriptional coregulator that fine-tunes target gene transcription by binding to specific nuclear components receptors (Zhang et al., 2024). The high correlation suggests that the *Nrip3* may represent a candidate upstream modulator associated with *Slc6a3* expression and even the broader *Slc6a3-Cadps2* co-expression network module.

**Figure 4.**
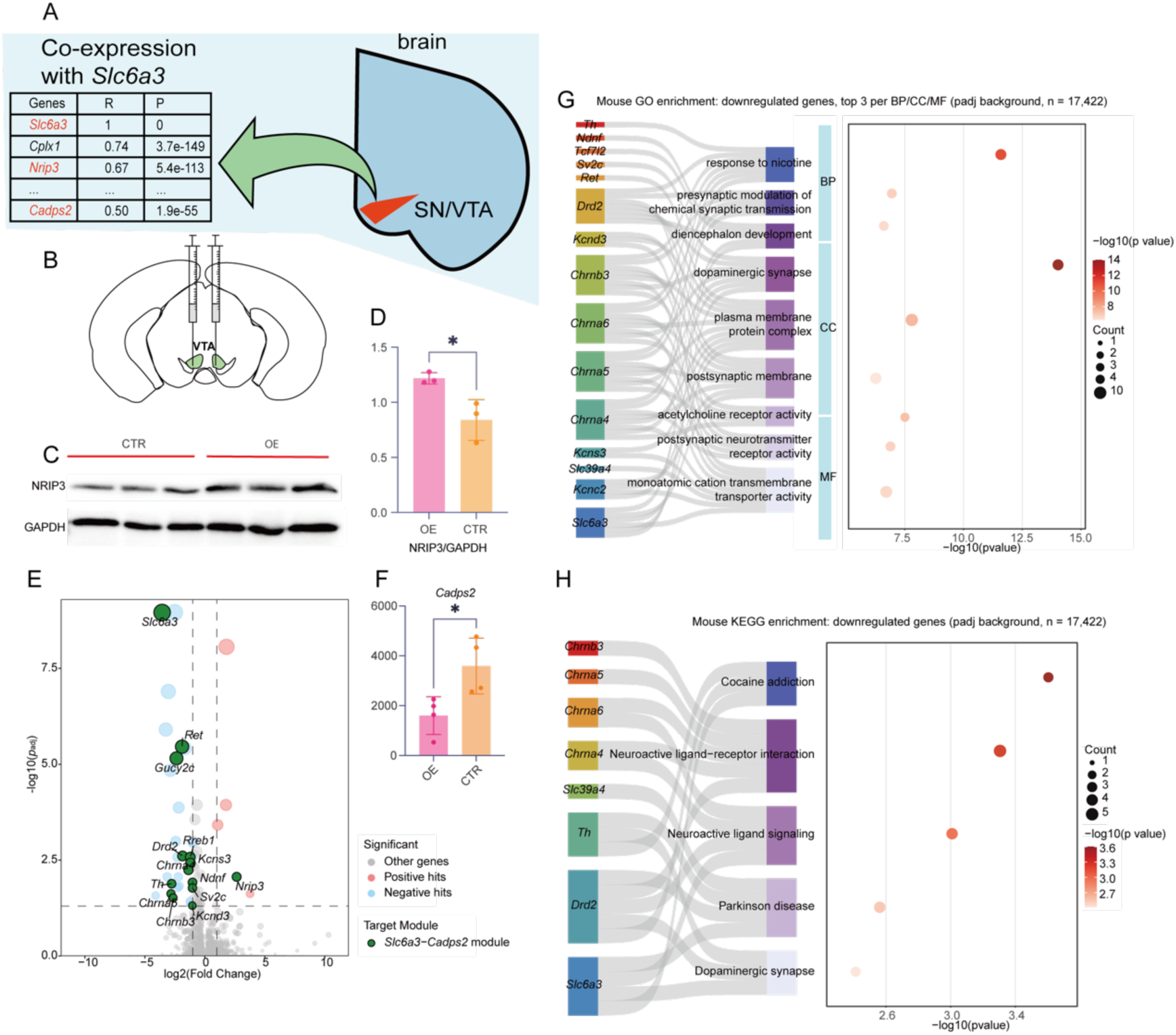
*Nrip3* overexpression in the ventral tegmental area (VTA) downregulates the *Slc6a3*-*Cadps2* co-expression network and alters dopamine-related transcriptional profiles. (A) *Slc6a3*-anchored spatial expression correlation landscape in the mouse midbrain. The analysis reveals highly synchronous co-expression of *Cadps2* and the transcriptional coregulator *Nrip3* with *Slc6a3* across cell populations in these regions. (B) Schematic representation of the experimental design. Recombinant adeno-associated viruses (rAAV-hSyn-mNrip3-P2A-EGFP for overexpression or rAAV-hSyn-EGFP for control) were injected bilaterally into the VTA of C57BL/6J mice (*n* = 15 per group). (C) Result of western blot of NRIP3 protein levels in the VTA. (D) Quantification of NRIP3 protein expression. NRIP3 levels in the overexpression group were approximately 44.5% higher than controls (unpaired *t*-test, *p* = 0.027, *n* = 3, biological replicates per group; The effect size was large, η² = 0.74). (E) Differential expression analysis (DESeq2, *p*.adj < 0.05 and |log2FC| > 1) from bulk RNA-seq of VTA tissues (*n* = 4 per group, biological replicates per group). There were 35 differentially expressed genes (DEGs) identified, of which 5 were significantly upregulated (including *Nrip3*) and 30 significantly downregulated. Notably, *Slc6a3* was the most significantly downregulated gene (log2FC = −3.56, *p*.adj = 1.10 × 10⁻⁹). (F) Targeted Mann-Whitney U test of *Cadps2* mRNA expression (two-tailed, *p* = 0.0286; median: control = 3522, overexpression = 1809), showing a significant decrease despite *Cadps2* not reaching the genome-wide DEG threshold (*p*.adj = 0.24). (G) GO enrichment analysis of the identified DEGs. A Sankey-bubble plot illustrates the significant enrichment observed in key functional terms, including response to nicotine, regulation of membrane potential, synaptic membrane, and monoatomic ion channel activity, with *Drd2*, *Slc6a3*, and associated channel/receptor genes identified as core functional genes within these networks. (H) KEGG pathway enrichment analysis of the identified DEGs. Significant enrichment was observed in pathways including cocaine addiction, neuroactive ligand-receptor interaction, Parkinson’s disease, and dopaminergic synapse, with *Drd2* and *Slc6a3* identified as primary functional driver genes within these pathways.

To further validate these correlations at the single-cell level while controlling for cell-type composition, we computed pairwise correlations among *Slc6a3*, *Cadps2*, and *Nrip3* within the annotated dopaminergic neuron subpopulation (*n* = 3,353 cells) using raw gene-level count matrix from the 10Xv3 single-cell RNA-seq dataset before normalization or log transformation. All three gene pairs showed significant positive correlations (*Slc6a3–Cadps2*: Pearson r = 0.65, Spearman ρ = 0.76; *Slc6a3–Nrip3*: Pearson r = 0.91, Spearman ρ = 0.80; *Cadps2–Nrip3*: Pearson r = 0.52, Spearman ρ = 0.60; all *p* < 10⁻²⁰⁰; Supplementary Figure S6; S8), confirming that the co-expression relationships among these three genes are not merely byproducts of dopaminergic neuron abundance variation.

To verify the regulatory role of *Nrip3* in the expression of *Slc6a3* and the *Slc6a3*-*Cadps2* co-expression network module, we mediated expression of *Nrip3* gene in the VTA via stereotactic injection of recombinant adeno-associated viruses (rAAVs). We bilaterally injected either an overexpression construct (rAAV-hSyn-mNrip3-P2A-EGFP; *n*=15) or an empty vector control (rAAV-hSyn-EGFP; *n*=15) into the VTA of C57BL/6J mice (Figure 4B). After 21 days of stable expression and behavioral tests finished, VTA tissues from 4 randomly selected mice per group were analyzed by bulk RNA sequencing (RNA-seq). Six samples (3 per group, each from an independent mouse) were prepared for western blotting (WB) and confirmed successful overexpression of the NRIP3 protein (unpaired *t*-test, *p* = 0.027, *n* = 3 biological replicates per group; Figure 4C & 4D). Differential expression analysis was performed using DESeq2 (*Nrip3*-overexpression vs. control, *n* = 4 biological replicates per group). DESeq2 size-factor normalization and dispersion estimation were applied; no LFC shrinkage or batch-effect correction was performed, as the eight samples were sequenced in a single batch. Genes with Benjamini-Hochberg-adjusted *p* (*p*.adj) < 0.05 and |log2FoldChange| > 1 were called as DEGs, yielding 35 DEGs (5 upregulated, 30 downregulated; Supplementary Table S10). *Slc6a3* was the most significantly downregulated gene (log2FoldChange = −3.56, lfcSE = 0.475, Wald stat = −7.49, *p*= 6.66 × 10⁻¹⁴, *p*.adj = 1.10 × 10⁻⁹; ∼11.8-fold downregulation) (Figure 4E). Of the 30 downregulated genes, 13 were derived from the the previously identified green module containing *Slc6a3* and *Cadps2*. These findings suggest that *Nrip3* may regulate the *Slc6a3-Cadps2* network module, potentially modulating the expression levels of multiple genes within it. Notably, while *Cadps2* showed a substantial fold change (log2FoldChange = −1.19), it did not pass the threshold in DESeq2 (*p*.adj = 0.24, above the 0.05 cutoff), likely due to the limited sample size (*n* = 4 per group). Given that *Cadps2* was a pre-specified candidate based on GWAS and co-expression network evidence, a hypothesis-driven targeted Mann-Whitney U test was performed on DESeq2 normalized counts (*p* = 0.0286), supporting its significant downregulation. This targeted test is explicitly labeled as post-hoc and is interpreted cautiously; it is not treated as genome-wide DEG evidence (Figure 4F). We conducted a GO enrichment analysis on the aforementioned DEGs, revealing their association with the response to nicotine, regulation of membrane potential, synaptic membrane, and dopaminergic synapse terms, which were significantly enriched. The enrichment in these biological processes and cellular components is predominantly driven by the *Drd2*, *Slc6a3*, and cholinergic receptor (*Chrna4/5/6*) genes (Figure 4G). Subsequently, we performed KEGG pathway enrichment analysis on the identified DEGs, which revealed significant enrichment in pathways associated with cocaine addiction, neuroactive ligand-receptor interactions, Parkinson’s disease, and dopaminergic synapses. The primary functional drivers within these pathways are the *Drd2* and *Slc6a3* genes (Figure 4H).

### Overexpression of *Nrip3* in the VTA induces behavior change in mice

To explore the influence of the *Nrip3* gene on behavior, we subjected the mice to a battery of tests (Figure 5A). Initial assessment involved a 3D kinematic analysis, which recorded the mice’s posture and locomotor activities within a circular open field apparatus over a 1-hour period (Figure 5B). We applied uniform manifold approximation and projection (UMAP) to reduce the dimensionality of 40 extracted kinematic features into a two-dimensional space. This analysis showed robust intra-group homogeneity within both the control and overexpression cohorts, with minimal sample dispersion. Notably, the kinematic profiles of all 30 samples mapped onto a continuous and curved trajectory within the low-dimensional manifold. Despite this continuity, distinct inter-group segregation was evident. These mice can be clustered into three relatively discrete subgroups, highlighting specific differences in motor performance between the control and overexpression groups, with only a minority of admixed samples (Figure 5C). Furthermore, individual classification based on these kinematic parameters yielded high accuracy (Figure 5D). Subsequent statistical comparison of kinematic parameters revealed that the overexpression mice exhibited a hyperactive phenotype characterized by increased locomotor activity, movement intensity, and postural extension. Specifically, the overexpression group displayed significantly elevated speed, movement intensity, and total distance traveled compared to controls (Figure 5E-G).

**Figure 5.**
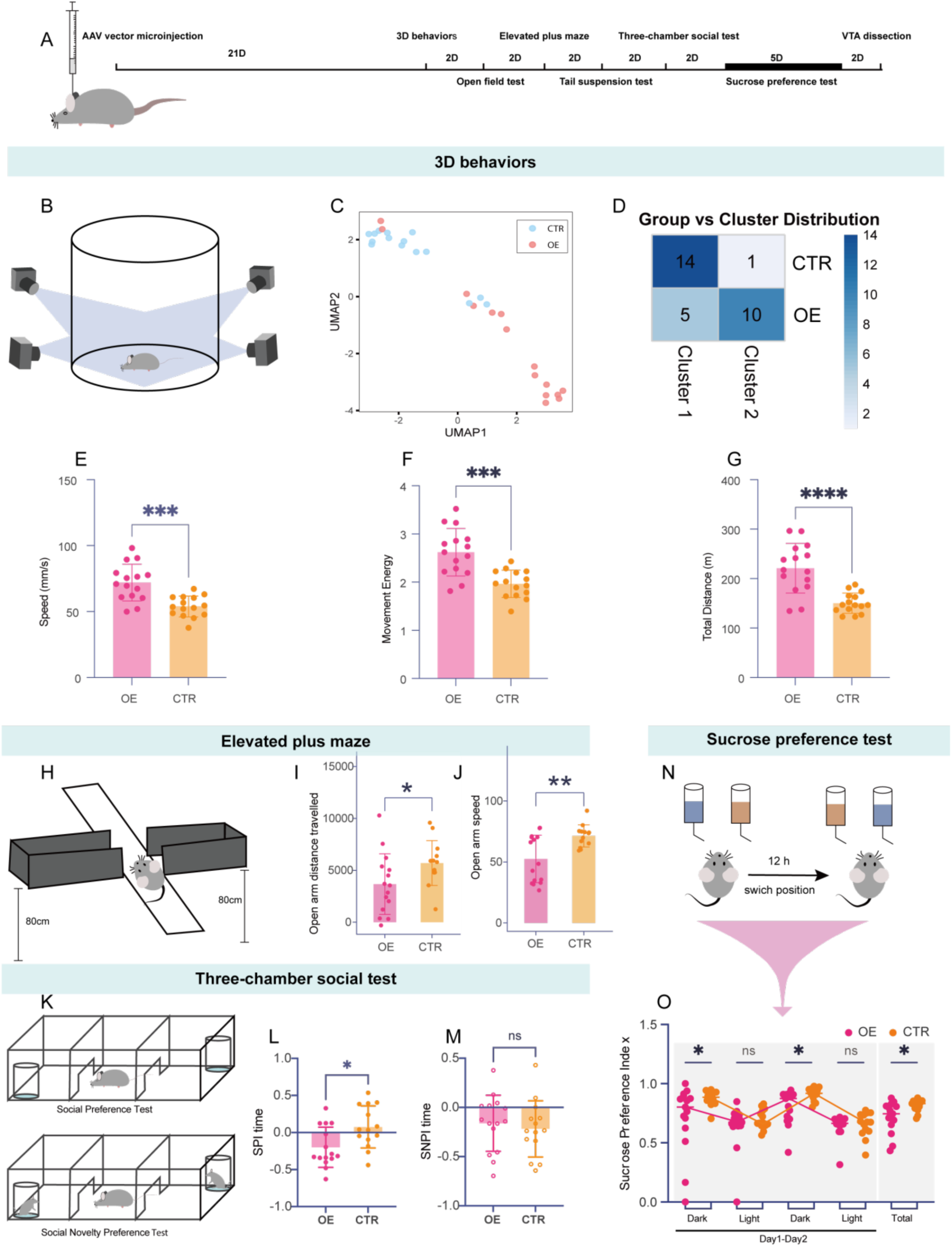
VTA-specific overexpression of *Nrip3* alters locomotor kinematics, open-arm exploration, social approach, and anhedonia in mice. (A) Timeline of behavioral experiments. (B–D) High-dimensional kinematic profiling via 3D behavioral testing. (B) Schematic of the 3D behavior capture system. (C) Uniform manifold approximation and projection (UMAP) visualization of 40 kinematic variables reduced to 2D space. Each dot represents one mouse sample (OE, *n* = 15; CTR, *n* = 15). CTR: control group; OE: *Nrip3*-overexpression group.. CTR: control group; OE: *Nrip3*-overexpression group. (D) Classification accuracy of individual mice based on locomotor kinematic parameters. (E–G) Statistical analysis of locomotor parameters (OE, *n* = 15; CTR, *n* = 15). *Nrip3*-overexpression mice showed increased movement speed (E; Welch’s t = 4.370, *P* = 0.0002), movement energy (F; Welch’s t = 4.474, *p* = 0.0002), and total distance traveled (G; Welch’s *t* = 5.064, *p* < 0.0001) compared to controls. (H–J) elevated plus maze (EPM) based open-arm exploration assessment. (H) Schematic of the EPM test. After adjustment for open filed test (OFT) speed, *Nrip3*-overexpression mice showed reduced open-arm distance travelled (I; OE, *n* = 15; CTR, *n* = 13; Welch’s *t* = 2.103, *p* = 0.0455) and reduced open-arm speed (J; OE, *n* = 15; CTR, *n* = 13; Welch’s *t* = 3.363, *p* = 0.0030). (K–M) Three-chamber social interaction test. (K) Schematic of the three-chamber social test. Social preference index (L) and social novelty preference index (M) during the social approach phase (OE, *n* = 15; CTR, *n* = 14). *Nrip3*-overexpression mice showed reduced social preference time index (L; Welch’s t = 2.652, *p* = 0.0133), whereas the social preference entry index was not significantly different between groups (M; Welch’s t = 0.410, *P* = 0.6853). (N) Schematic of the sucrose preference test (SPT). (O) Sucrose preference was measured across four consecutive 12-h light/dark intervals and across the overall test period (OE, *n* = 15; CTR, n = 14). *Nrip3*-overexpression mice showed reduced sucrose preference during the first dark phase (Welch’s *t* = 2.184, *p* = 0.0447), the second dark phase (Welch’s *t* = 2.835, *p* = 0.0110), and the overall test period (Welch’s *t* = 2.702, *p* = 0.0145), whereas differences during the light phases were not significant (first light phase: *p* = 0.2928; second light phase: *p* = 0.8095). Note: Data are presented as mean ± SEM. Statistical significance was determined using two-tailed unpaired *t* tests with Welch’s correction. For locomotion-sensitive OFT and EPM endpoints, OFT speed was included as a locomotor covariate, and group comparisons were performed on speed-adjusted individual values. Detailed statistical reporting for mouse behavioral endpoints is provided in Supplementary Table S7. \**p* < 0.05, \*\**p* < 0.01, \*\*\**p* < 0.001.

To evaluate anxiety-related behavior in these mice, we performed the open field test (OFT) and the elevated plus maze (EPM) (Figure 5H). Because *Nrip3*-overexpression mice exhibited altered baseline locomotor activity in the 3D behavioral assay, we further evaluated locomotion-sensitive OFT and EPM endpoints after adjustment for OFT speed. After this covariate adjustment, OFT immobility time, center-zone time, and center-zone entries remained non-significant between groups (Supplementary Figure S9). In the EPM test, open-arm time and open-arm entries also remained non-significant (Supplementary Figure S10), whereas open-arm distance travelled and open-arm speed remained significantly lower in the *Nrip3*-overexpression group (Figure 5I, J). These results suggest reduced open-arm exploration after accounting for baseline locomotor differences and may reflect a change in an anxiety-related behavioral dimension. However, because canonical anxiety indices in the OFT and EPM, including center-zone measures, open-arm time, and open-arm entries, were not consistently altered, this interpretation should remain cautious.

To evaluate social preference and interaction motivation, we employed the three-chamber social test (Figure 5K). During the social approach phase, *Nrip3* overexpression mice showed a significantly reduced social preference time index, whereas the social preference entry index did not differ significantly between groups (Figure 5L, M). In the social novelty phase, neither the social novelty preference time index nor the entry index differed significantly between groups (Supplementary Figure S11). These findings indicate that *Nrip3* upregulation severely impairs proactive social motivation, leading to social withdrawal.

Given the central role of the VTA dopaminergic system in reward and emotional processing (Morales & Margolis, 2017), we used the tail suspension test (TST) and sucrose preference test (SPT) to evaluate depressive-like states and reward sensitivity. TST results showed no significant difference in total struggling time or the count of struggles between the two groups (Supplementary Figure S12). However, in the SPT (Figure 5N), *Nrip3* overexpressing mice showed a significantly reduced overall sucrose preference. Notably, When analyzed across four consecutive 12-hour intervals, this reduction was evident during the dark phases but not during the light phases (Figure 5O). These findings suggest a phase-dependent reduction in sucrose preference in *Nrip3*-overexpression mice. The difference observed during the dark phase may be due to the nocturnal activity pattern of mice.

## Discussion

In this study, we investigated genetic architecture underlying stranger-directed social anxiety-like behavior in CKD, a breed officially recognized for its role in police and security operations in China. We identified a series of genes including *FAIM2, ASIC1, SLC6A3,* and *CADPS2* as candidate genes. Notably, most of these genes are known to be associated with human psychiatric disorders. *FAIM2* encodes a neuroprotective apoptotic inhibitor that has been widely studied as a regulator of neuronal survival and cellular stress responses (Planells-Ferrer et al., 2016). It plays a role in psychiatric disorders, such as major depressive disorder, and its variants have been robustly linked to metabolic traits (Littleton et al., 2024). *ASIC1* encodes an acid-sensing ion channel that is abundantly expressed in the amygdala and plays a prominent role in anxiety-related behavior. It plays critical role in innate fear and panic in mouse models (Pidoplichko et al., 2014; Ziemann et al., 2009). *SLC6A3* encodes the human dopamine transporter, which plays a role in maintaining monoaminergic balance, a vital factor for proper synaptic transmission. *SLC6A3* dysfunctions have been linked to ADHD, substance use disorders, and severe mood dysregulation in human studies (M. E. Reith et al., 2022). It has been reported that DAT knockdown can lead to reduced anxiety in mice (M. E. A. Reith et al., 2022). *CADPS2* regulates the calcium-dependent exocytosis of dense-core vesicles and is involved in neurotrophin release. It is also crucial for normal synaptic connectivity. *CADPS2* deficiencies have been linked to severe depressive-like behaviours and autism spectrum disorders (Tetsushi Sadakata et al., 2007). The identification of these genes associated with SAD-like behavior in dogs implies shared genetics between canine anxiety and human psychiatric disorders. Importantly, the social anxiety-like phenotype in CKDs was stranger-specific, affected dogs displayed abnormal behaviors (trembling, freezing, avoidance, tail-tucking) exclusively toward unfamiliar strangers, while showing normal behavior toward familiar handlers and trainers. This stranger-specificity supports the characterization of the observed phenotype as social anxiety-like and aligns with a core feature of social anxiety disorder. However, because our experimental paradigm specifically assessed anxiety-related responses toward unfamiliar humans, we cannot rule out the possibility that these dogs may also exhibit generalized fearfulness, neophobia, reduced sociability, or other anxiety-related traits in other contexts.

By integrating whole-brain spatial and single-cell transcriptomic data from mice, we further revealed that *Slc6a3* and *Cadps2* may not function in isolation; rather, they belong to a highly synergistic co-expression module localized in the VTA and SNc. This finding supports the biological plausibility that dysregulation of this co-expression module may constitute to anxiety or fear-related behavior, although direct validation in CKDs is still required. Moreover, this study innovatively integrated GWAS of domestic dogs with co-expression network analysis in mice. This strategy provides a novel analytical framework for GWAS with small-scale samples. This approach can be used to detect suggestive signals and minor-effect loci that are typically overlooked in conventional GWAS-only analyses.

Integrating spatial single cell and bulk transcriptomic analysis, we showed a regulatory association of *Nrip3* within the *Slc6a3-Cadps2* co-expression network, which also involves *Slc18a2* and *Drd2*. We overexpressed the *Nrip3* gene in the VTA of mice. Behavioral results showed altered locomotor activity, reduced social approach, reduced sucrose preference, and reduced EPM open-arm exploration in mice. The EPM open-arm distance and speed findings may indicate a change in a specific anxiety-related behavioral dimension, but this interpretation should remain cautious because conventional anxiety indices, including OFT center-zone measures and EPM open-arm time and entries, were not consistently altered. The difference between OFT and EPM findings may reflect the fact that these assays probe partially overlapping but non-identical behavioral dimensions: OFT center-zone measures mainly capture exploration and center avoidance in a novel open arena, whereas EPM open-arm measures additionally involve height exposure, open-arm risk assessment, and conflict between exploration and avoidance. Thus, the mouse behavioral data are best interpreted as broad behavioral alterations involving activity, social approach, reward-related behavior, and possible anxiety-related open-arm exploration, rather than definitive evidence of a generalized anxiety-like phenotype. NRIP3, a nuclear receptor-interacting protein, likely depends on interactions with the nuclear receptor to perform its functions. In the midbrain dopamine system, the only known nuclear receptor is nuclear receptor subfamily 4 group A member 2 (NR4A2/NURR1), which shows selectively high expression in the substantia nigra pars compacta/ventral tegmental area (SNc/VTA). Simultaneously, NR4A2 has established roles in dopaminergic neuron development and in the transcriptional regulation of downstream target genes such as *Slc6a3* and *Drd2*. (Bruning et al., 2019; Kamath et al., 2022a). Consistent with this possible route, UCSC JASPAR2026 TFBS annotations also identified NR4A2 motif sites within both the human *SLC6A3* locus and the mouse *Slc6a3* locus. Therefore, NRIP3 may affect *Slc6a3* expression partly through NR4A2-associated regulatory mechanisms. However, this remains a mechanistic hypothesis, and direct experimental validation will be required. Furthermore, our analyses revealed that rAAV-mediated overexpression of *Nrip3* led to downregulation of *Slc6a3* expression in this experimental context. This observation appears to contradict the positive correlation observed between *Nrip3* and *Slc6a3* in single-cell transcriptomic datasets. This discrepancy remains an unresolved mechanistic issue. One plausible but unproven hypothesis is that the sophisticated homeostatic regulation of the dopaminergic system (Best et al., 2009) may trigger negative feedback-like loops that could mask or reverse the canonical regulatory relationship observed under physiological, steady-state conditions. Additionally, rAAV-mediated delivery, particularly when utilizing strong constitutive promoters, often results in supra-physiological protein levels (Van Alstyne et al., 2021) that may exceed the operational range of the neuron’s native transcriptional network, potentially triggering compensatory regulatory feedback absent under physiological conditions. Further studies are needed to resolve this contradictory regulation. Several limitations of the *Nrip3* overexpression intervention should be noted. First, the intervention was performed in adult mice, which may not model the congenital developmental processes wherein pathogenic genes exhibit abnormal expression from embryonic or early postnatal stages. Second, the viral overexpression approach resulted in supra-physiological *Nrip3* levels, which may not reflect the effects of natural regulatory variation. Third, the hSyn promoter drives expression in all neurons rather than being specific to dopaminergic neurons, limiting the ability to attribute the observed effects specifically to the dopaminergic system. Fourth, only gain-of-function (overexpression) was performed; loss-of-function experiments (e.g., shRNA, CRISPRi, or conditional knockout using Dat-Cre mice) were attempted but did not achieve sufficient viral intervention efficacy and were excluded, and remain a priority for future work. The canine causal chain, from genetic variation through *SLC6A3-CADPS2* network to *Nrip3*-mediated regulation to behavior, is therefore inferred rather than directly demonstrated. Future studies should employ Dat-Cre conditional, developmental stage-specific, and physiological-level manipulations (e.g., CRISPRa), as well as loss-of-function rescue experiments, to establish causality. Future studies should test this proposed route using co-immunoprecipitation to examine potential NRIP3-NR4A2 interaction, ChIP-qPCR or CUT&Tag/CUT&RUN to assess NR4A2-associated occupancy at candidate *Slc6a3* regulatory regions, and luciferase reporter assays to evaluate whether these regulatory sequences respond to NRIP3 and/or NR4A2 manipulation.

Although this study proposes a highly promising neurogenetic mechanism, several objective limitations remain. First, constrained by the strict early selection and elimination mechanisms for police dogs, individuals harboring social anxiety-like behavior are largely eliminated during the puppy stage. This extremely low natural retention rate resulted in a relatively limited sample size of affected dogs (18 cases) during the GWAS phase of this study, representing the full set of available affected CKD individuals within the study period. The absence of an independent replication cohort further limits the generalizability of the GWAS findings, which should be regarded as preliminary candidate signals. Additionally, while population structure and sex were included as covariates, other potential confounders (age, kennel, lineage, training history, and housing conditions) could not be fully incorporated due to incomplete records and may introduce residual confounding. Regarding the canine behavioral phenotype, several methodological limitations should be acknowledged. Initially, the behavioral assessment was retrospective and semi-structured; blinding of evaluators to the dogs’ prior behavioral history could not be fully confirmed. Subsequently, inter-rater reliability analysis could not be performed due to the retrospective nature of the assessment and the limited availability of video recordings. Furthermore, each dog was assessed only once, precluding evaluation of test-retest stability across different days or strangers. Finally, the phenotype classification used qualitative judgment rather than a standardized scoring scale. These limitations are acknowledged, and future studies should employ prospective, standardized behavioral protocols with blinded scoring, inter-rater reliability metrics (e.g., Cohen’s κ), and test-retest validation. A CKD breeding colony with social anxiety-like individuals has been established to support such future work. Additionally, while we controlled for cell-type composition by computing the *Slc6a3–Cadps2* correlation within the dopaminergic neuron subpopulation using raw read counts (Pearson r = 0.65, Spearman ρ = 0.76), the bulk WGCNA co-expression (r = 0.96) partly reflects tissue-level cell-type composition, and direct functional interaction between *Slc6a3* and *Cadps2* remains to be validated through conditional perturbation experiments. Second, given the multiple constraints of high costs, technical barriers, and stringent ethical reviews associated with *in vivo* gene editing in large mammals, our current mechanistic validation primarily relied on the mouse model. While we verified the function of this network at the murine level, we cannot complete a causal closed-loop demonstration within the CKD cohort. The mouse *Nrip3* overexpression experiment provides correlational evidence from a gain-of-function intervention, but does not establish that canine genetic variants cause disease through the dopamine system. Although the evidence from GWAS, co-expression analysis, *in silico* perturbation, and mouse overexpression experiments converges on a coherent candidate mechanism, these findings do not constitute complete causal proof. Therefore, this study should be interpreted as hypothesis-generating, and causal validation will require further experimental work. Given the significant genetic and evolutionary divergence between species, and even among distinct dog breeds, it is essential to account for potential, non-negligible disparities in underlying pathological pathways when extrapolating findings across models. This urgently requires future efforts that leverage canine-derived neural *in vitro* models, construct single-cell brain transcriptomic atlases of both diseased and healthy dogs, or introduce more advanced *in vivo* research systems for dogs to achieve final validation.

Finally, the viral intervention performed in adult mouse brains in this study may not perfectly model the congenital pathological developmental processes wherein pathogenic genes exhibit abnormal expression from the embryonic or early postnatal stages. Simultaneously, the inherent physical spatial deviations of stereotaxic injections, along with the local inflammatory and immune responses triggered by invasive procedures, may potentially confound precise behavioral evaluations. To more precisely dissect the physiological functions of this dopaminergic network, future experimental paradigms should employ conditional gene mutation or knockout models utilizing the Cre/loxP system (e.g., *Dat-Cre* mice). Such approaches would enable the cell-type-specific manipulation of network components, thereby providing a more rigorous framework for establishing a direct causal link between gene expression, dopaminergic signaling, and specific behavioral outcomes.

## Materials and Methods

### General Information

#### Study Design and Analysis Plan

The canine GWAS was designed a confirmatory analysis to identify loci associated with social anxiety-like behavior in CKDs. The downstream analyses-including WGCNA, scTenifoldKnk in silico perturbation, gsMap spatial correlation, *Nrip3* upstream inference, and enrichment analyses-were exploratory. Throughout the manuscript, confirmatory analyses use the term “tested,” while exploratory analyses use “explored” or “prioritized.” The primary outcome of the GWAS was genome-wide significant loci (*SLC6A3* on chr34, chr27 locus); suggestive loci (*CADPS2* on chr14) were considered secondary. For the mouse experiments, the primary endpoints were EPM, OFT, three-chamber social test metrics, sucrose preference; secondary endpoints included 3D kinematic parameters, and TST immobility. This study was not pre-registered, but the analysis plan was internally established before implementation, and no analyses were conducted based on post-hoc inspection of results. Sample size for the GWAS (18 cases, 82 controls) was determined by the full set of available affected CKD individuals within the study period, constrained by early police-dog selection. Sample size for mouse experiments (*n* = 15 per group) was based on standard practices in behavioral neuroscience.

#### Software and Parameters

A comprehensive summary of all software tools and versions used in this study is provided in Supplementary Table S8: BaseNumber (v1.0.4.0), GEMMA (v0.98.5), PLINK (v1.9), Scanpy (v1.10.2), scTenifoldKnk (v1.0.0), WGCNA (v1.72-5), clusterProfiler (v4.4.3, R 4.4.3), STAR (v2.7.10a), featureCounts (v2.0.8), fastp (v0.23.2), DESeq2 (v1.46.0, R 4.3.0), GraphPad Prism (v10), Cytoscape (v3.10.4), STRING (v12.0), gsMap (v1.72.3), and the BehaviorAtlas 3D-AI system. Versions are also cited at first mention in the respective Methods subsections.

### Behavior Phenotyping and Sample Collection in the Chinese Kunming Dogs

In this study, venous blood samples were collected from a cohort of 103 CKDs for GWAS analysis. Following genomic quality control, three individuals were excluded as heterozygosity outliers (genome-wide heterozygosity rate deviating more than ±3 standard deviations from the population mean, detected using PLINK --het), yielding a final cohort of 100 CKDs. Concurrent with blood sampling, basic demographic information (e.g., age, sex, and baseline fearfulness status) was recorded, and a phenotypic assessment of social behavior was conducted (Ethics approval number: IACUC-OE-2023-02-001). Dogs were evaluated in an outdoor environment while being restrained on a leash by their caretakers, with an unfamiliar person approaching the dogs to assess their behavioral responses. During the testing phase, experimenters evaluated whether the dogs exhibited overt behavioral traits indicative of social anxiety when confronted by a stranger, such as trembling, freezing, and avoidance,, tail-tucking, or urination. Each dog was assessed once; no repeated tests on different days or with different strangers were conducted, Multiple observers concurrently evaluated each dog. Dogs were classified as cases (social anxiety-like) if they exhibited marked trembling, freezing, avoidance, tail-tucking, or urination when approached by the unfamiliar stranger and classified as controls if they displayed proactive approach and interactive behaviors toward the stranger. Classification was based on qualitative judgment (presence/absence of overt behavioral signs) rather than a quantitative scoring threshold, determined by consensus among multiple evaluators. Notably, case dogs displayed abnormal behaviors only toward the unfamiliar stranger; when familiar individuals (handlers or trainers) were present, no such behaviors were observed, supporting the stranger-specificity of the phenotype, but not excluding the possibility of generalized fearfulness or neophobia. Behavior phenotyping and blood sample collection were predominantly conducted at the Kunming Police Dog Base of the Ministry of Public Security, with additional samples obtained from dogs kept in affiliated kennel under the base’s supervision.

### Whole-genome sequencing data and GWAS

#### Variant Calling

The collected blood samples were directly sent to Novogene Co., Ltd. (Beijing, China) for WGS utilizing the DNBSEQ-T7 high-throughput sequencing platform. The raw sequencing data underwent rigorous quality control (QC) processing to trim low-quality reads and adapter sequences, ensuring data accuracy and reliability. The cleaned sequencing data were then aligned to the dog reference genome (CanFam4) using the BaseNumber DNA Sequencing Data Analysis Software, developed by SaiLe Gene. The BaseNumber software suite is an ultra-fast, proprietary bioinformatics pipeline (Auwera & O’Connor, 2020; DePristo et al., 2011; McKenna et al., 2010; Zhang et al., 2021). By implementing highly parallelized algorithms optimized for a heterogeneous “CPU + GPU” computing environment, these tools substantially accelerate the efficiency of DNA sequencing data analysis. The analytical workflows are constructed by chaining various tools via a workflow system, primarily encompassing germline and somatic variant-calling pipelines (including GATK Mutect2- and VarDict-based algorithms). While ensuring high concordance of results, the BaseNumber pipeline dramatically reduces computational time and overhead. Currently, this software can process diverse data types generated by next-generation sequencing (NGS) platforms, including WGS, whole-exome sequencing (WES), and targeted panels, and has broad applications across genetic disease research, oncology, population cohort studies, and agricultural genomics. The alignment process mapped short sequencing reads to the reference sequence, generating alignment files (BAM format) that were subsequently processed for duplicate removal and alignment optimization. Following this, the SLC (SaiLe Caller) module in the BaseNumber software was used to perform variant calling on the aligned data, identifying SNPs and InDels at a genome-wide scale. The SLC module yields outputs in both VCF and gVCF formats, with the latter (gVCF) utilized for subsequent joint variant analysis. Finally, the slmgvcf_gpu program was deployed to merge the gVCF files of all individual samples into a single, comprehensive VCF file. The genetic variants captured within this merged VCF file constituted the core genotypic dataset for the downstream GWAS.

#### Data Quality Control

The merged VCF file was converted into PLINK format for rigorous quality control using PLINK v1.9 (Chang et al., 2015). The stringent QC criteria applied were as follows: (1) exclusion of individuals with a missing genotype rate > 2%; (2) exclusion of single-nucleotide polymorphisms (SNPs) with a missing rate > 2%; (3) removal of SNPs with a minor allele frequency (MAF) < 0.15. This specific MAF threshold was empirically optimized among 0.05, 0.10, 0.15, and 0.20 to maximize the retention of robust independent signals while effectively mitigating false-positive artifacts in our specific sample size; (4) removal of SNPs exhibiting extreme deviation from Hardy-Weinberg equilibrium (HWE) with a *P* < 1 × 10^−4^; and (5) exclusion of outlier individuals whose genome-wide heterozygosity rate deviated by more than ± 3 standard deviations (SD) from the population mean, thereby circumventing potential biases arising from sample contamination or extreme inbreeding.

#### Genome-Wide Association Analysis

To account for potential confounding effects arising from population structure and cryptic relatedness, the GWAS was conducted using the LMM implemented in GEMMA (Zhou & Stephens, 2012). Specifically, we first computed a VanRaden standardized genetic relationship matrix (GRM) using the - gk option to quantify genotypic similarity among all individuals and estimate the covariance structure. Population structure was assessed by principal component analysis (PCA) using 174,290 LD-pruned (r²<0.2) variants, and the first two principal components (PC1 and PC2) together with sex were included as fixed-effect covariates, and the first two principal components (PC1 and PC2) together with sex were included as fixed-effect covariates. Association tests were performed using the -lmm 4 option, which employs a restricted maximum likelihood algorithm to accurately estimate both fixed and random effects, with the GRM as the random-effect component. We note that the GEMMA LMM outputs beta coefficients (β) with standard errors (SE) rather than odds ratios; effect sizes (β, SE, and 95% CI) for the lead SNPs are reported in Supplementary Table S4. Other potential confounders (age, kennel, lineage, training history, and housing conditions) could not be fully incorporated due to incomplete records and are discussed as a limitation.

To strictly control the family-wise error rate for multiple testing across the genome, a genome-wide significance threshold was determined using the Bonferroni correction. Based on the number of relatively independent SNPs retained after linkage disequilibrium (LD) pruning (a total of 1,426,148 SNPs, r²<0.7), the strict significance threshold was set at *p <* 3.51×10^−8^. Furthermore, a suggestive threshold of *p <* 1×10^−5^ was additionally adopted, following the convention used in human GWAS.

### Mouse Brain Multi-Transcriptomic Analysis

Processing of mouse midbrain scRNA-seq Data. The midbrain single-cell transcriptomic data were sourced from a publicly available adult mouse whole-brain cell atlas. To characterize cellular heterogeneity and gene profiles within the VTA, we used midbrain scRNA-seq data generated via the 10x Genomics v3 platform from this study (Yao et al., 2023).

#### *In Silico* Gene Knockout Simulation

Standard preprocessing of the midbrain scRNA-seq data, including quality control, doublet removal, clustering, and cell type annotation, was performed using Scanpy (Wolf et al., 2018). Dopaminergic neurons were annotated based on a canonical marker set including *Th*, *Drd2*, *Slc6a3*, and *Nr4a2*, and were isolated to extract the raw count matrix for subsequent analyses. Subsequently, the scTenifoldKnk (Osorio et al., 2022) was employed to conduct *in silico* knockouts of *Slc6a3* and *Cadps2*. A pipeline-specific stringent QC was applied to the input matrix (mitochondrial ratio < 0.1, minimum library size = 1000, minimum gene expression fraction = 0.001). A robust wild-type single-cell gene regulatory network (WT scGRN) was constructed via principal component regression and tensor decomposition, reinforced by random subsampling (nc_nNet = 10, nc_nCells = 500, nc_nComp = 3).

*In silico* knockouts simulated by setting all regulatory edge weights linked to the target gene to zero, creating a pseudo-KO scGRN. Non-linear manifold alignment then projected both WT and pseudo-KO networks into a shared latent space. Downstream gene perturbation significance was evaluated using Euclidean distance, Box-Cox Z-scores, and FDR-adjusted *p*. Significantly perturbed genes were subjected to GO and KEGG enrichment analyses to delineate the biological networks and mechanisms underlying the target genes.

### Mouse Whole-Brain Spatial Transcriptomic Data

We analyzed previously published spatial transcriptomic datasets of the adult mouse whole brain. Following preprocessing, these datasets were stored as h5ad files, each containing a gene expression count matrix, cell type annotations, and the spatial coordinates of the spots (Chen et al., 2022).

#### gsMap Analysis

gsMap is a computational tool leveraging graph neural networks (GNNs) to integrate spatial coordinates with gene expression profiles in spatial transcriptomics. It identifies homogeneous regions for individual spots and evaluates whether trait-associated genetic variants are enriched in the vicinity of spatially specific genes. To achieve a more robust evaluation of the spatial specificity of gene expression, we employed gsMap to compute the gene spatial specificity scores (GSSs) (Song et al., 2025). The resultant GSS values for each spot or cell were subsequently mapped back to the corresponding spatial transcriptomic datasets. The generated feather files were utilized for subsequent gene expression correlation analyses.

#### Preliminary Identification of the *Slc6a3-Cadps2* Co-expression Network Module

Single-cell spatial transcriptomic data are inherently plagued by a high proportion of technical zeros (dropouts), resulting in substantial missingness in gene expression detection. Such data sparsity directly compromises the accurate estimation of inter-genic correlations and introduces biases into the construction of co-expression networks. To circumvent this limitation, we extracted information specifically for dopaminergic neurons within the SN/VTA from the feather files generated via gsMap processing of the mouse whole-brain spatial transcriptomic data. These files encapsulate the GSS for every gene within each individual cell (Song et al., 2025).

Subsequently, using the GSSs, we plotted the whole-brain spatial expression profiles of the target genes to compare the expression patterns of *Slc6a3* and *Cadps2* across distinct brain regions. Concurrently, both Pearson and Spearman correlation coefficients were calculated between the GSS of each globally expressed gene and that of *Slc6a3*. Genes were then ranked by the magnitude of these correlation coefficients to generate a comprehensive ranked list of *Slc6a3* gene expression correlations.

#### Identification of the *Slc6a3-Cadps2* Co-expression Module via WGCNA

Bulk RNA-seq data from the VTA were obtained from GEO accession GSE228031 (Browne et al., 2023). We used the GEO-provided processed CPM expression matrix (GSE228031_deseq2_VTA_cpm.csv.gz) rather than raw FASTQ files. Gene identifiers were mapped to gene symbols; genes without valid symbols were removed. CPM values were log2-transformed as log2(CPM + 1), low-expression genes were filtered (rowMeans > 1), and the top 25% most variable genes were retained (n = 3,432). Samples were further normalized using limma normalizeBetweenArrays, and quality control was performed using the WGCNA goodSamplesGenes function, yielding 38 samples after outlier removal. The WGCNA (Langfelder & Horvath, 2008) was employed to construct the co-expression network with the following parameters: soft-thresholding power = 13 (scale-free topology fit R² = 0.858), minimum module size = 100, deepSplit = 2, and merge cut height = 0.5. A gene co-expression similarity matrix was generated by calculating Pearson correlation coefficients across all gene pairs. To robustly approximate a scale-free topology network (R^2^ > 0.8), an optimal soft-thresholding power (β) was applied to transform the similarity matrix into a weighted adjacency matrix. This matrix was subsequently converted into a topological overlap matrix (TOM) to capture intricate, indirect genic interactions and mitigate background noise. Finally, average linkage hierarchical clustering was performed based on the TOM-based dissimilarity measure (1 - TOM), and the dynamic tree cut algorithm was utilized to automatically delineate independent gene modules exhibiting highly synergistic expression patterns.

#### Cell-Type-Controlled Correlation Analysis

To control for potential confounding by cell-type composition in bulk tissue, pairwise correlations among *Slc6a3*, *Cadps2*, and *Nrip3* were computed within the annotated dopaminergic neuron subpopulation (n = 3,353 cells) using raw counts from single-cell RNA-seq data, where cell type is held constant. Both Pearson and Spearman correlations were calculated.

#### Protein-Protein Interaction Network Construction and Hub Gene Screening

To elucidate the interactions among the proteins encoded by the target genes, the filtered mouse target gene list was uploaded to the STRING online database (version 12.0) (Szklarczyk et al., 2023). The organism was set to *Mus musculus*. By integrating interaction data from various sources including experimental validation, co-expression, and text mining, the minimum required interaction score threshold was set to 0.400 (medium confidence), and isolated nodes without any connections were excluded. The constructed PPI network was imported into Cytoscape software (version 3.10.4) (Shannon et al., 2003) for topological analysis. In the network visualization, the node size, node color gradient, and edge thickness were mapped to the nodes’ degree centrality (Degree). Additionally, the cytoHubba plugin in Cytoscape was used to calculate node topological scores using the Degree algorithm, thereby identifying the top-ranked hub genes with the highest connectivity, facilitating the subsequent discovery of key molecular targets.

### Viral-Mediated Gene Intervention in the Mouse Model

#### Experimental Animals

Thirty C57BL/6J male mice (6–7 weeks old, weighing 17–21 g) were purchased from SPF (Beijing) Biotechnology Co., Ltd. The mice were housed, with five animals per cage, in a conventional facility at 21°C under a 12:12 h light-dark cycle, with lights on at 08:00 and lights off at 20:00. After a one-week acclimation period, stereotaxic brain injections were performed (Ethics approval number: IACUC-RE-2026-02-007).

#### Stereotaxic Injection of Adeno-Associated Virus (AAV)

Adeno-associated virus serotype 9 (AAV9) vectors were synthesized by Brain Case (Wuhan) Biotechnology Co., Ltd. For the gain-of-function experiment, a total of 2 viral vectors were constructed which are the rAAV-hSyn-mNrip3-P2A-EGFP and rAAV-hSyn-EGFP plasmids. Transcript of *Nrip3* is NM_020610.1. Following a one-week acclimation period, mice underwent stereotaxic surgery. Anesthesia was maintained with isoflurane (0.75% concentration, 0.5 L/min gas flow). The anesthetized mice were then secured onto a stereotaxic frame. Based on *The Mouse Brain in Stereotaxic Coordinates* (2nd Edition) [67] and optimized through preliminary experiments, bilateral injections targeting the VTA (Coordinates: anteroposterior [AP]: −2.92 mm, mediolateral (Noyvert et al.): ± 0.45 mm, dorsoventral (Boeva et al.): −4.4 mm) were performed using a microsyringe connected to a microinjection pump. A total viral volume of 200 nL per side was delivered at a constant rate of 0.5 nL per second. The needle was left in place for an additional 8 minutes post-injection to facilitate optimal viral diffusion and prevent backflow along the needle tract.

Subsequently, the microsyringe was slowly withdrawn, and the scalp incision was closed with sutures. Post-operatively, the mice were placed in a temperature-controlled warming chamber to recover, preventing fatal hypothermia. Throughout the surgical procedure, the isoflurane concentration was continuously monitored and dynamically adjusted to ensure adequate depth of anesthesia while preventing fatal anesthetic overdose. Behavioral experiments were conducted 21 days post-injection to ensure sufficient time for robust viral gene expression.

#### Attempted *Nrip3* knockdown experiment

To explore the loss-of-function effects of *Nrip3*, we also attempted an AAV-mediated knockdown experiment using an shRNA strategy targeting the longest mouse *Nrip3* transcript (NM_020610.1; Gene ID: 78593). The selected construct was packaged as an AAV9 vector, rAAV-hSyn-EGFP-5’miR-30a-shRNA(mNrip3)-3’miR-30a, with a measured titer of 5.01 × 10¹² vg/mL. The virus was delivered by stereotaxic injection using the same VTA-targeting coordinates and injection procedure described above. Knockdown efficacy was evaluated by measuring *Nrip3* expression in the injected brain region. Under the present experimental conditions, AAV9-shRNA(mNrip3) injection did not produce a reliable reduction in *Nrip3* expression and therefore did not constitute an effective loss-of-function perturbation. These data were therefore not included as interpretable loss-of-function evidence in the main analysis, but this technical outcome may provide a useful reference for future AAV-mediated *Nrip3* knockdown studies.

### Mouse Behavioral Assays

#### General behavioral testing and tracking conditions

Behavioral tests were conducted during the light phase between 08:00 and 18:00, whereas the sucrose preference test was continuously monitored across both light and dark phases over two days. Behavioral assays were performed under dim ambient illumination; however, the exact illuminance level was not quantitatively recorded. Before each behavioral assay, mice were transferred to the testing room and allowed to habituate for 30–60 min. Open field, elevated plus maze, tail suspension, and three-chamber social behavior were recorded and analyzed using VisuTrack 3.0 behavioral tracking software (Shanghai XinRuan Information Technology Co., Ltd., Shanghai, China). Three-dimensional kinematic behavior was recorded and analyzed using the 3D-AI Mouse Behavior Analysis System (BehaviorAtlas: https://bayonesci.com/behavioratlas/) (Huang et al., 2021).

#### 3D Behavioral Analysis

As previously described (Cai et al., 2026), 3D motion capture and spontaneous behavior decomposition were conducted to quantitatively evaluate rodent behavior. We employed the 3D motion capture and behavioral decomposition techniques based on the hierarchical 3D motion learning framework proposed by Huang *et al* (Huang et al., 2021). This pipeline consists of three primary steps: synchronous multi-view video acquisition, 3D posture reconstruction, and the subsequent decomposition of behavioral dynamics from skeletal trajectories.

These operations were executed using the BehaviorAtlas, which integrates hardware and software components to enable automated multi-camera synchronization, robust 3D skeleton tracking, and machine-learning-based behavioral profiling. Four cameras positioned at the corners of the apparatus synchronously recorded the spontaneous behavior of the mice for 60 minutes. The BehaviorAtlas system tracked and analyzed the movement trajectories of 16 anatomical keypoints on the mouse (including the nose, ears, neck, back, paws, limbs, and the base, middle, and tip of the tail). This tracking ultimately identified 40 distinct behavioral clusters, which were subsequently manually categorized into basic behavioral phenotypes, including sniffing, grooming, rearing, jumping, turning, and locomotion. The proportion of each behavior was calculated by dividing the duration of the specific behavior by the total duration of all recorded behaviors.

#### Open Field Test (OFT)

Prior to testing, mice were transferred to the behavioral testing room and habituated to the environment for 30 to 60 minutes. The inner and outer surfaces of the open field apparatus (50 cm × 50 cm × 35 cm) were wiped down with 75% ethanol to remove odors and residues. The arena was virtually divided into a 3 × 3 grid, further defining three functional zones: the central zone, the peripheral zone, and the corners. Each mouse was gently placed in the center of the arena and allowed to explore freely. Locomotion trajectories were continuously recorded for 10 minutes using VisuTrack 3.0 behavioral tracking software.. Between each trial, the apparatus was immediately cleared of any feces and thoroughly wiped with 75% ethanol to eliminate olfactory cues, ensuring no interference with the subsequent animal’s behavior. For data analysis, the time spent in and the number of entries into the different zones during the first 5 minutes were extracted as core metrics to evaluate anxiety-like behaviors (Tye et al., 2011).

#### Elevated Plus Maze (EPM)

Prior to testing, mice were transferred to the testing room for a 30 – 60 minutes habituation period. The maze apparatus was thoroughly cleaned with 75% ethanol before each trial to remove any residual odors or traces from the previous animal. VisuTrack 3.0 behavioral tracking software was calibrated to define the maze into three distinct regions: the open arms, closed arms, and center area. Each mouse was gently picked up and placed in the central area facing an open arm, with its back to the experimenter. The experimenter then swiftly and quietly exited the testing area. The software automatically recorded the movement trajectory of the mouse for 5 minutes. After the trial, the mouse was gently returned to its home cage, and the apparatus was thoroughly cleaned with 75% ethanol and paper towels, then prepared for the next subject. Anxiety-like behaviors were assessed by calculating the time spent in and the number of entries into the open arms, closed arms, and the center area (Walf & Frye, 2007).

#### Tail Suspension Test (TST)

The experiment was conducted in a quiet environment. Each mouse was gently removed from its home cage, and medical adhesive tape was applied approximately 1 cm from the tip of the tail. The mouse was then suspended downwards from the suspension apparatus, ensuring the snout was positioned approximately 30 cm above the floor. Behavior was recorded for 6 minutes and analyzed using VisuTrack 3.0 behavioral tracking software. The cumulative immobility time during the last 4 minutes of the test was extracted as the primary evaluation metric. Prolonged immobility is generally indicative of more pronounced depression-like behavior. Following the test, the tape was gently removed, the mouse was returned to its home cage, and all experimental equipment was thoroughly wiped and disinfected (Tye et al., 2013).

#### Three-Chamber Social Test

The three-chamber social test was conducted to evaluate sociability and social memory/recognition (Gunaydin et al., 2014). Prior to testing, mice were habituated to the testing room for 30 – 60 minutes. To eliminate olfactory cues, the entire apparatus and enclosures were thoroughly cleaned with 75% ethanol between all trials. The experimental procedure consisted of three consecutive 10-minute phases. In the first phase (habituation), the test mouse was placed in the center chamber with the doors open, allowing it to freely explore all three empty chambers to acclimate to the spatial environment. In the second phase (sociability), an unfamiliar, sex-matched conspecific mouse (Stranger 1) was enclosed in a transparent, perforated cage in one side chamber, while an identical empty cage was placed in the opposite chamber. The test mouse was allowed to freely explore, enabling assessment of social preference between the social object and the empty cage. In the third phase (social novelty), a novel, unfamiliar mouse (Stranger 2) was introduced into the previously empty cage, alongside the now-familiar Stranger 1. The test mouse was granted another 10 minutes of free exploration to assess social recognition and memory. For data recording, VisuTrack 3.0 behavioral tracking software was used to continuously record the trials. The time spent in each chamber, the number of entries, and the duration of active interaction with the stranger mice were extracted as key metrics to evaluate social tendencies and recognition capabilities.

#### Sucrose Preference Test (SPT)

As previously described (Yang et al., 2025), mice were individually housed and habituated to two bottles of water for 2 days, followed by a 24-hour water-deprivation period prior to testing. During the test, mice were provided with two bottles: one containing regular water and the other containing a 1% sucrose solution. The acute sucrose preference test was conducted during the first 3 hours of the mice’s active phase. To eliminate spatial preference bias, the positions of the two bottles were alternated every 0.5 hours. Water and sucrose consumption were accurately recorded.

The sucrose preference rate was defined as the percentage of sucrose solution consumed relative to the total liquid consumption over the 3-hour period. In the chronic sucrose preference test, sucrose consumption was continuously monitored over the subsequent 2 days. Bottle positions were alternated during the light phase, and liquid consumption was measured every 12 hours to ensure measurement balance across different spatial locations and temporal periods.

#### Behavioral Data Analysis

Mice were randomly assigned to their respective treatment groups. All behavioral assays were conducted with anonymized subjects, and the experimenters remained strictly blinded to the specific treatment conditions of the animals. No data from any surviving individuals were excluded from the analysis. Statistical analyses were performed using GraphPad Prism 10 software. Two-group comparisons of behavioral measurements were performed using unpaired t tests with Welch’s correction. For behavioral endpoints potentially confounded by baseline locomotor activity, an ANCOVA-style covariate adjustment was performed using OFT speed as the locomotor covariate. For each endpoint, a linear model was fitted as outcome ∼ group + OFT speed, and locomotor-speed-adjusted individual values were calculated as: adjusted value = observed value − β × (OFT speed − mean OFT speed), where β represents the common slope estimated from the model. Group differences in these adjusted values were then assessed using two-tailed unpaired t tests with Welch’s correction. Sucrose preference measurements was also compared between groups using unpaired t tests with Welch’s correction. Specific sample sizes (*n*, representing individual animals) are detailed in the corresponding figure legends, and detailed statistical reporting for mouse behavioral endpoints, including exact *p* values, confidence intervals, effect-size estimates, and FDR-adjusted *p* values, is provided in Supplementary Table S7_. Data are presented as the mean ± standard error of the mean (SEM) or as individual data points.

### RNA-seq Analysis of the VTA

#### Sample Collection

Following the completion of the behavioral assays, a total of 8 mice (*n* = 4 for the overexpression group and *n* = 4 for the control group) were euthanized. The left ventral tegmental area (VTA) was rapidly microdissected. The isolated tissue samples were immediately sent to Novogene Co., Ltd. (Beijing, China) for transcriptomic sequencing (RNA-seq).

#### Raw Read Quality Assessment and Filtering

Raw sequencing quality was assessed using metrics provided by Novogene. Across the eight samples, with four biological replicates per group, the number of raw reads ranged from 40.5 to 45.7 million. The Q30 and Q20 rates ranged from 97.95% to 98.37% and from 99.19% to 99.34%, respectively. GC content ranged from 48.7% to 51.0%, and the sequencing error rate was 0.01% for all samples. Following quality filtering with fastp, 96.6%–98.6% of the raw reads were retained as clean reads.

#### Transcriptomic Data Processing and Quantification

Raw paired-end FASTQ data were processed with fastp (v0.23.2) (Chen et al., 2018) to remove adapter sequences and trim low-quality bases. The cleaned reads were then aligned to the mouse reference genome (GRCm38.p6) using STAR (Dobin et al., 2013), with the --sjdbOverhang parameter set to 149 based on the corresponding GTF annotation file. Subsequently, gene-level expression quantification was performed using featureCounts from the Subread package (v2.0.8) (Liao et al., 2014). The quantification was executed with 8 threads (-T 8) in paired-end mode (-p) to ensure accurate fragment counting. Finally, an R script (version 4.3.0) was used to merge the individual sample-count files into a unified raw gene expression count matrix (merged_counts.txt) for downstream statistical analyses.

#### Differential Expression Analysis

Differential expression analysis was performed using DESeq2 (v1.46.0) in R 4.3.0, using the merged raw count matrix (merged_counts.txt) produced by featureCounts as input. The model was design = ∼group, comparing the *Nrip3*-overexpression group against the control (n = 4 per group); no additional covariates were included. DESeq2 size-factor normalization and dispersion estimation were applied; no LFC shrinkage or batch-effect correction (e.g., SVA, ComBat) was performed, as the eight samples were sequenced in a single batch. Genes with Benjamini-Hochberg-adjusted *p* (padj) < 0.05 and |log2FoldChange| > 1 were called as DEGs. For Cadps2, which did not pass the genome-wide DEG threshold (padj = 0.24), a hypothesis-driven targeted Mann-Whitney U test was performed on DESeq2 normalized counts (two-tailed, *p*= 0.0286), explicitly labeled as post-hoc. The full, unfiltered DEG table, including both significant and non-significant genes tested in the DESeq2 analysis, is provided in Supplementary Table S8.

#### Gene Enrichment Analysis

GO and KEGG pathway enrichment analyses were performed using the clusterProfiler R package (version 4.4.3) with the org.Mm.eg.db annotation database. Target mouse gene symbols were mapped and converted to Entrez IDs. The statistical background was defined separately for each enrichment analysis according to the gene set entering the corresponding analysis. For the scTenifoldKnk virtual perturbation analyses shown in Figure 2, the statistical background was defined as the shared set of genes evaluated in both the *Slc6a3* and *Cadps2* virtual perturbation outputs. Specifically, we intersected the gene lists from the two scTenifoldKnk differential regulation tables and obtained 2,680 mouse genes; this shared tested-gene set was used as the background universe for overlap statistics and was mapped to mouse Entrez IDs for GO and KEGG enrichment analyses. For the WGCNA/PPI candidate-gene enrichment analysis, the universe was defined from the mouse VTA expression dataset GSE228031 after gene-symbol processing, removal of duplicated symbols, and low-expression filtering using rowMeans > 1, resulting in 13,726 expressed background genes, of which 13,566 were successfully mapped to mouse Entrez IDs. GO and KEGG enrichment analyses were then performed using these analysis-specific backgrounds rather than the default whole-genome background. GO enrichment was performed across three domains: biological process (BP), cellular component (CC), and molecular function (MF), with significance defined as nominal *p* < 0.05. KEGG pathway enrichment used the same universe and significance threshold. For the overlap analysis between *Slc6a3* and *Cadps2* perturbed gene sets, significance was assessed using Fisher’s exact test with the same 2,680-gene universe, using genes significant at *p*.adj < 0.05 as input. All enrichment analysis results were visualized using the ggplot2 R package. For the *Nrip3*-overexpression RNA-seq enrichment analysis, the enrichment background was defined as the 17,422 genes with non-missing adjusted *p* values in the DESeq2 analysis. Among these, 16,951 genes were successfully mapped to mouse Entrez IDs and used as the effective GO/KEGG universe. This background was chosen because the foreground downregulated genes were selected from the same DESeq2 testing framework using *p*.adj < 0.05 and log2FoldChange < −1.

### Western blot

Target cells were lysed in radioimmunoprecipitation assay (RIPA) buffer (Thermo Scientific) on ice for 30 min, the cell lysates were centrifuged at 14,000g for 15 min at 4 °C, and the supernatants were collected. Protein concentration was determined using the BCA protein assay kit (Thermo Scientific). Briefly, 20ul protein liquid was mixed with 160 ul BCA Working fluid for 30 min at 37 °C, and the absorbance was measured at 562 nm with Microplate Reader (BIO-TEK), protein concentrations were calculated from a standard curve. For immunoblotting, 20 µg proteins from each sample were separated by 10% SDS–polyacrylamide gel using Mini-PROTEAN Tetra (Bio-Rad) and then transferred to PVDF membrane (Millipore) using Trans-Blot system (Bio-Rad). The PVDF membranes were blocked with 5% bovine serum albumin (BSA) (beyotime) in TBS/T for 2 h at room temperature, followed by incubation with primary antibodies overnight at 4 °C: Nrip3 (1:1000, HUABIO, Cat. No. ER64225) and GAPDH (1:1000, Abclonal, Cat. No. AC001). After washing, the membranes were incubated with second antibodies for 1 h at room temperature: Anti-rabbit IgG, HRP-linked Antibody (1:1000, CST, Cat. No. 7074S). The membranes were incubated with BeyoECL Moon (beyotime) for 1-2 min, signals were visualized by ChemiDoc XRS+ System (Bio-Rad).

## ACKNOWLEDGEMENTS

We thank the Animal Bank at the Germplasm Bank of Wild Species (https://cstr.cn/31121.02.GBOWS.AB) for providing biological materials and technical support.

## DATA AND CODE AVAILABILITY

The raw whole-genome sequencing data and RNA-seq data produced in this study are archived in the Genome Sequence Archive (GSA) at the National Genomics Data Center (NGDC) under the accession number PRJCA062986. All processed datasets and corresponding analysis scripts (including GWAS, WGCNA, scTenifoldKnk *in silico* knockout, enrichment analysis, and differential expression analysis) have been deposited at Zenodo (DOI: 10.5281/zenodo.21816555). All data and code are permanently archived and citable via DOI.

## Reference

Alex, E., Gennotte, P., Morros Nuevo, A., Yu, Y., Keep, B., Sullivan, M., Mills, D., Warrier, V., & Raffan, E. (2025). GWAS for behavioral traits in golden retrievers identifies genes implicated in human temperament, mental health, and cognition. Proceedings of the National Academy of Sciences, 122(48), e2421757122.

Auwera, G.A., & O’Connor, B.D. (2020). Genomics in the cloud: using Docker, GATK, and WDL in Terra. (*No Title)*.

Baba, A., Kloiber, S., & Zai, G. (2022). Genetics of social anxiety disorder: a systematic review. Psychiatric Genetics, 32(2). https://journals.lww.com/psychgenetics/fulltext/2022/04000/genetics_of_social_anxie ty_disorder a_systematic.1.aspx

Bahi, A., & Dreyer, J.-L. (2019). Dopamine transporter (DAT) knockdown in the nucleus accumbens improves anxiety-and depression-related behaviors in adult mice. Behavioural Brain Research, 359, 104–115.

Bandyopadhyay Prasanta, S., Forster Malcolm, R., Oxford, E., Barkow Jerome, H., Leda, C., John, T., William, B., Richardson Robert, C., Beck Aaron, T., & John, R.A. (2014). American Psychiatric Association, Diagnostic and Statistical Manual of Mental Disorders: Dsm-5, Washington, DC, American Psychiatric Publishing, 2013. Ananth Mahesh, In defense of an evolutionary concept of health nature, norms, and human biology, Aldershot, England, Ashgate. Philosophy, 39(6), 683–724.

Best, J.A., Nijhout, H.F., & Reed, M.C. (2009). Homeostatic mechanisms in dopamine synthesis and release: a mathematical model. Theoretical Biology and Medical Modelling, 6(1), 21. 10.1186/1742-4682-6-21

Boeva, V., Louis-Brennetot, C., Peltier, A., Durand, S., Pierre-Eugène, C., Raynal, V., Etchevers, H.C., Thomas, S., Lermine, A., Daudigeos-Dubus, E., et al. (2017). Heterogeneity of neuroblastoma cell identity defined by transcriptional circuitries. Nature Genetics, 49(9), 1408–1413. 10.1038/ng.3921

Bonora, E., Graziano, C., Minopoli, F., Bacchelli, E., Magini, P., Diquigiovanni, C., Lomartire, S., Bianco, F., Vargiolu, M., & Parchi, P. (2014). Maternally inherited genetic variants of CADPS2 are present in autism spectrum disorders and intellectual disability patients. EMBO Molecular Medicine, 6(6), 795–809.

Browne, C.J., Futamura, R., Minier-Toribio, A., Hicks, E.M., Ramakrishnan, A., Martínez-Rivera, F.J., Estill, M., Godino, A., Parise, E.M., & Torres-Berrío, A. (2023). Transcriptional signatures of heroin intake and relapse throughout the brain reward circuitry in male mice. Science Advances, 9(23), eadg8558.

Bruning, J.M., Wang, Y., Oltrabella, F., Tian, B., Kholodar, S.A., Liu, H., Bhattacharya, P., Guo, S., Holton, J.M., & Fletterick, R.J. (2019). Covalent modification and regulation of the nuclear receptor Nurr1 by a dopamine metabolite. Cell chemical biology, 26(5), 674–685. e676.

Cai, X., Sun, T., Feng, M., Chen, G., Zhou, J., Zhuang, H., Wang, D., Chen, Y., Cheng, Z., & Xu, Z. (2026). Taurocholic acid is associated with disturbed functional connectivity in the hippocampus of patients with depression. Advanced Science, 13(13), e08693.

Chang, C.C., Chow, C.C., Tellier, L.C., Vattikuti, S., Purcell, S.M., & Lee, J.J. (2015). Second-generation PLINK: rising to the challenge of larger and richer datasets. Gigascience, 4(1), s13742–13015-10047-13748.

Chen, A., Liao, S., Cheng, M., Ma, K., Wu, L., Lai, Y., Qiu, X., Yang, J., Xu, J., & Hao, S. (2022). Spatiotemporal transcriptomic atlas of mouse organogenesis using DNA nanoball-patterned arrays. Cell, 185(10), 1777–1792. e1721.

Chen, S., Zhou, Y., Chen, Y., & Gu, J. (2018). fastp: an ultra-fast all-in-one FASTQ preprocessor. Bioinformatics, 34(17), i884–i890.

Demin, K.A., Krotova, N.A., Ilyin, N.P., Galstyan, D.S., Kolesnikova, T.O., Strekalova, T., de Abreu, M.S., Petersen, E.V., Zabegalov, K.N., & Kalueff, A.V. (2022). Evolutionarily conserved gene expression patterns for affective disorders revealed using cross-species brain transcriptomic analyses in humans, rats and zebrafish. Scientific Reports, 12(1), 20836. 10.1038/s41598-022-22688-x

DePristo, M.A., Banks, E., Poplin, R., Garimella, K.V., Maguire, J.R., Hartl, C., Philippakis, A.A., Del Angel, G., Rivas, M.A., & Hanna, M. (2011). A framework for variation discovery and genotyping using next-generation DNA sequencing data. Nature Genetics, 43(5), 491–498.

Dobin, A., Davis, C.A., Schlesinger, F., Drenkow, J., Zaleski, C., Jha, S., Batut, P., Chaisson, M., & Gingeras, T.R. (2013). STAR: ultrafast universal RNA-seq aligner. Bioinformatics, 29(1), 15–21.

Donner, J., Freyer, J., Davison, S., Anderson, H., Blades, M., Honkanen, L., Inman, L., Brookhart-Knox, C.A., Louviere, A., & Forman, O.P. (2023). Genetic prevalence and clinical relevance of canine Mendelian disease variants in over one million dogs. PLoS Genetics, 19(2), e1010651.

Dutrow, E.V., Serpell, J.A., & Ostrander, E.A. (2022). Domestic dog lineages reveal genetic drivers of behavioral diversification. Cell, 185(25), 4737–4755. e4718.

Fiorenzano, A., Storm, P., Sozzi, E., Bruzelius, A., Corsi, S., Kajtez, J., Mudannayake, J., Nelander, J., Mattsson, B., & Åkerblom, M. (2024). TARGET-seq: Linking single-cell transcriptomics of human dopaminergic neurons with their target specificity. Proceedings of the National Academy of Sciences, 121(47), e2410331121.

Fujima, S., Yamaga, R., Minami, H., Mizuno, S., Shinoda, Y., Sadakata, T., Abe, M., Sakimura, K., Sano, Y., & Furuichi, T. (2021). CAPS2 Deficiency Impairs the Release of the Social Peptide Oxytocin, as Well as Oxytocin-Associated Social Behavior. J Neurosci, 41(20), 4524–4535. 10.1523/jneurosci.3240-20.2021

Grison, S., Paul, M.A., Kessler, K., & Tipper, S.P. (2005). Inhibition of object identity in inhibition of return: Implications for encoding and retrieving inhibitory processes. Psychonomic Bulletin & Review, 12(3), 553–558.

Gunaydin, L.A., Grosenick, L., Finkelstein, J.C., Kauvar, I.V., Fenno, L.E., Adhikari, A., Lammel, S., Mirzabekov, J.J., Airan, R.D., & Zalocusky, K.A. (2014). Natural neural projection dynamics underlying social behavior. Cell, 157(7), 1535–1551.

Halo, J.V., Pendleton, A.L., Shen, F., Doucet, A.J., Derrien, T., Hitte, C., Kirby, L.E., Myers, B., Sliwerska, E., & Emery, S. (2021). Long-read assembly of a Great Dane genome highlights the contribution of GC-rich sequence and mobile elements to canine genomes. Proceedings of the National Academy of Sciences, 118(11), e2016274118.

Han, M., Chang, J., & Kim, J. (2016). Loss of divalent metal transporter 1 function promotes brain copper accumulation and increases impulsivity. Journal of neurochemistry, 138(6), 918–928.

Hong, C.J., Yeon, J., Yeo, B.K., Woo, H., An, H.K., Heo, W., Kim, K., & Yu, S.W. (2020). Fas-apoptotic inhibitory molecule 2 localizes to the lysosome and facilitates autophagosome-lysosome fusion through the LC3 interaction region motif–dependent interaction with LC3. The FASEB Journal, 34(1), 161–179.

Huang, K., Han, Y., Chen, K., Pan, H., Zhao, G., Yi, W., Li, X., Liu, S., Wei, P., & Wang, L. (2021). A hierarchical 3D-motion learning framework for animal spontaneous behavior mapping. Nat. Commun. 12, 2784. In.

Huang, Q.-G., Zhang, M.-Z., Zhang, S.-J., Ge, X.-Z., Fu, J., Wen, T.-G., Peng, J.-G., Wang, G.-D., Wang, S.-Z., & Wang, L. (2024). Genetic study on heritability and novel SNP loci of temperament in Chinese Kunming Dog. Reproduction and Breeding, 4(1), 1–4.

Iguchi, H., Katsuzawa, T., Saruta, C., Sadakata, T., Kobayashi, S., Sato, Y., Sato, A., Sano, Y., Maezawa, S., & Shinoda, Y. (2024). Calcium-dependent activator protein for secretion 2 is involved in dopamine release in mouse midbrain neurons. Frontiers in Molecular Neuroscience, 17, 1444629.

Iguchi, H., Katsuzawa, T., Saruta, C., Sadakata, T., Kobayashi, S., Sato, Y., Sato, A., Sano, Y., Maezawa, S., Shinoda, Y., et al. (2024). Calcium-dependent activator protein for secretion 2 is involved in dopamine release in mouse midbrain neurons [Brief Research Report]. Frontiers in Molecular Neuroscience, Volume 17 - 2024. 10.3389/fnmol.2024.1444629

Johannesen, K.M., Liu, Y., Koko, M., Gjerulfsen, C.E., Sonnenberg, L., Schubert, J., Fenger, C.D., Eltokhi, A., Rannap, M., Koch, N.A., et al. (2022). Genotype-phenotype correlations in SCN8A-related disorders reveal prognostic and therapeutic implications. Brain, 145(9), 2991–3009. 10.1093/brain/awab321

Kamath, T., Abdulraouf, A., Burris, S., Langlieb, J., Gazestani, V., Nadaf, N.M., Balderrama, K., Vanderburg, C., & Macosko, E.Z. (2022a). Single-cell genomic profiling of human dopamine neurons identifies a population that selectively degenerates in Parkinson’s disease. Nature neuroscience, 25(5), 588–595.

Kamath, T., Abdulraouf, A., Burris, S.J., Langlieb, J., Gazestani, V., Nadaf, N.M., Balderrama, K., Vanderburg, C., & Macosko, E.Z. (2022b). Single-cell genomic profiling of human dopamine neurons identifies a population that selectively degenerates in Parkinson’s disease. Nature neuroscience, 25(5), 588–595. 10.1038/s41593-022-01061-1

Kreitmaier, P., Katsoula, G., & Zeggini, E. (2023). Insights from multi-omics integration in complex disease primary tissues. Trends in Genetics, 39(1), 46–58. 10.1016/j.tig.2022.08.005

Langfelder, P., & Horvath, S. (2008). WGCNA: an R package for weighted correlation network analysis. BMC bioinformatics, 9(1), 559.

Liao, Y., Smyth, G.K., & Shi, W. (2014). featureCounts: an efficient general purpose program for assigning sequence reads to genomic features. Bioinformatics, 30(7), 923–930.

Lindblad-Toh, K., Wade, C.M., Mikkelsen, T.S., Karlsson, E.K., Jaffe, D.B., Kamal, M., Clamp, M., Chang, J.L., Kulbokas III, E.J., & Zody, M.C. (2005). Genome sequence, comparative analysis and haplotype structure of the domestic dog. Nature, 438(7069), 803–819.

Littleton, S.H., Trang, K.B., Volpe, C.M., Cook, K., DeBruyne, N., Maguire, J.A., Weidekamp, M.A., Hodge, K.M., Boehm, K., & Lu, S. (2024). Variant-to-function analysis of the childhood obesity chr12q13 locus implicates rs7132908 as a causal variant within the 3′ UTR of FAIM2. Cell Genomics, 4(5).

McKenna, A., Hanna, M., Banks, E., Sivachenko, A., Cibulskis, K., Kernytsky, A., Garimella, K., Altshuler, D., Gabriel, S., & Daly, M. (2010). The Genome Analysis Toolkit: a MapReduce framework for analyzing next-generation DNA sequencing data. Genome research, 20(9), 1297–1303.

Meadows, J.R., Kidd, J.M., Wang, G.-D., Parker, H.G., Schall, P.Z., Bianchi, M., Christmas, M.J., Bougiouri, K., Buckley, R.M., & Hitte, C. (2023). Genome sequencing of 2000 canids by the Dog10K consortium advances the understanding of demography, genome function and architecture. Genome biology, 24(1), 187.

Miklósi, Á., & Topál, J. (2013). What does it take to become &#x2018;best friends&#x2019;? Evolutionary changes in canine social competence. Trends in Cognitive Sciences, 17(6), 287–294. 10.1016/j.tics.2013.04.005

Morales, M., & Margolis, E.B. (2017). Ventral tegmental area: cellular heterogeneity, connectivity and behaviour. Nature Reviews Neuroscience, 18(2), 73–85.

Morrill, K., Chen, F., & Karlsson, E. (2023). Comparative neurogenetics of dog behavior complements efforts towards human neuropsychiatric genetics. Human Genetics, 142(8), 1231–1246.

Nepal, B., Das, S., Reith, M.E., & Kortagere, S. (2023). Overview of the structure and function of the dopamine transporter and its protein interactions. Frontiers in physiology, 14, 1150355.

Noyvert, B., Erzurumluoglu, A.M., Drichel, D., Omland, S., Andlauer, T.F.M., Mueller, S., Sennels, L., Becker, C., Kantorovich, A., Bartholdy, B.A., et al. (2025). Imputation of structural variants using a multi-ancestry long-read sequencing panel enables identification of disease associations. In: eLife Sciences Publications, Ltd.

Osorio, D., Zhong, Y., Li, G., Xu, Q., Yang, Y., Tian, Y., Chapkin, R.S., Huang, J.Z., & Cai, J.J. (2022). scTenifoldKnk: An efficient virtual knockout tool for gene function predictions via single-cell gene regulatory network perturbation. Patterns, 3(3).

Overall, K.L. (2000). Natural animal models of human psychiatric conditions: assessment of mechanism and validity. Progress in Neuro-Psychopharmacology and Biological Psychiatry, 24(5), 727–776.

Paci, P., Fiscon, G., Conte, F., Wang, R.-S., Farina, L., & Loscalzo, J. (2021). Gene co-expression in the interactome: moving from correlation toward causation via an integrated approach to disease module discovery. npj Systems Biology and Applications, 7(1), 3. 10.1038/s41540-020-00168-0

Petri, A., Sullivan, A., Allen, K., & Sachs, B.D. (2024). Genetic loss of the dopamine transporter significantly impacts behavioral and molecular responses to sub-chronic stress in mice. Frontiers in Molecular Neuroscience, 17, 1315366.

Pidoplichko, V.I., Aroniadou-Anderjaska, V., Prager, E.M., Figueiredo, T.H., Almeida-Suhett, C.P., Miller, S.L., & Braga, M.F. (2014). ASIC1a activation enhances inhibition in the basolateral amygdala and reduces anxiety. Journal of Neuroscience, 34(9), 3130–3141.

Planells-Ferrer, L., Urresti, J., Coccia, E., Galenkamp, K.M., Calleja-Yagüe, I., López-Soriano, J., Carriba, P., Barneda-Zahonero, B., Segura, M.F., & Comella, J.X. (2016). Fas apoptosis inhibitory molecules: more than death-receptor antagonists in the nervous system. Journal of neurochemistry, 139(1), 11–21.

Prakash, N., Brodski, C., Naserke, T., Puelles, E., Gogoi, R., Hall, A., Panhuysen, M., Echevarria, D., Sussel, L., & Weisenhorn, D.M.V. (2006). A Wnt1-regulated genetic network controls the identity and fate of midbrain-dopaminergic progenitors in vivo.

Range, F., & Virányi, Z. (2015). Tracking the evolutionary origins of dog-human cooperation: the “Canine Cooperation Hypothesis” [Focused Review]. Frontiers in Psychology, Volume 5-2014. 10.3389/fpsyg.2014.01582

Reaume, C.J., & Sokolowski, M.B. (2011). Conservation of gene function in behaviour. Philosophical Transactions of the Royal Society B: Biological Sciences, 366(1574), 2100–2110. 10.1098/rstb.2011.0028

Reith, M.E., Kortagere, S., Wiers, C.E., Sun, H., Kurian, M.A., Galli, A., Volkow, N.D., & Lin, Z. (2022). The dopamine transporter gene SLC6A3: multidisease risks. Molecular psychiatry, 27(2), 1031–1046.

Reith, M.E.A., Kortagere, S., Wiers, C.E., Sun, H., Kurian, M.A., Galli, A., Volkow, N.D., & Lin, Z. (2022). The dopamine transporter gene SLC6A3: multidisease risks. Molecular Psychiatry, 27(2), 1031–1046. 10.1038/s41380-021-01341-5

Sadakata, T., Kakegawa, W., Mizoguchi, A., Washida, M., Katoh-Semba, R., Shutoh, F., Okamoto, T., Nakashima, H., Kimura, K., & Tanaka, M. (2007). Impaired cerebellar development and function in mice lacking CAPS2, a protein involved in neurotrophin release. Journal of Neuroscience, 27(10), 2472–2482.

Sadakata, T., Shinoda, Y., Oka, M., Sekine, Y., Sato, Y., Saruta, C., Miwa, H., Tanaka, M., Itohara, S., & Furuichi, T. (2012). Reduced axonal localization of a Caps2 splice variant impairs axonal release of BDNF and causes autistic-like behavior in mice. Proc Natl Acad Sci U S A, 109(51), 21104–21109. 10.1073/pnas.1210055109

Sadakata, T., Washida, M., Iwayama, Y., Shoji, S., Sato, Y., Ohkura, T., Katoh-Semba, R., Nakajima, M., Sekine, Y., Tanaka, M., et al. (2007). Autistic-like phenotypes in Cadps2-knockout mice and aberrant CADPS2 splicing in autistic patients. J Clin Invest, 117(4), 931–943. 10.1172/jci29031

Sarviaho, R., Hakosalo, O., Tiira, K., Sulkama, S., Salmela, E., Hytönen, M.K., Sillanpää, M.J., & Lohi, H. (2019). Two novel genomic regions associated with fearfulness in dogs overlap human neuropsychiatric loci. Translational Psychiatry, 9(1), 18. 10.1038/s41398-018-0361-x

Shannon, P., Markiel, A., Ozier, O., Baliga, N.S., Wang, J.T., Ramage, D., Amin, N., Schwikowski, B., & Ideker, T. (2003). Cytoscape: a software environment for integrated models of biomolecular interaction networks. Genome research, 13(11), 2498–2504.

Shi, P., Zhang, M.-J., Liu, A., Yang, C.-L., Yue, J.-Y., Hu, R., Mao, Y., Zhang, Z., Wang, W., & Jin, Y. (2023). Acid-sensing ion channel 1a in the central nucleus of the amygdala regulates anxiety-like behaviors in a mouse model of acute pain. Frontiers in Molecular Neuroscience, 15, 1006125.

Song, L., Chen, W., Hou, J., Guo, M., & Yang, J. (2025). Spatially resolved mapping of cells associated with human complex traits. Nature, 641(8064), 932–941.

Stein, M.B., Chen, C.-Y., Jain, S., Jensen, K.P., He, F., Heeringa, S.G., Kessler, R.C., Maihofer, A., Nock, M.K., Ripke, S., et al. (2017). Genetic risk variants for social anxiety. American Journal of Medical Genetics Part B: Neuropsychiatric Genetics, 174(2), 120–131. 10.1002/ajmg.b.32520

Suresh, H., Crow, M., Jorstad, N., Hodge, R., Lein, E., Dobin, A., Bakken, T., & Gillis, J. (2023). Comparative single-cell transcriptomic analysis of primate brains highlights human-specific regulatory evolution. Nature Ecology & Evolution, 7(11), 1930–1943. 10.1038/s41559-023-02186-7

Sutter, N.B., Eberle, M.A., Parker, H.G., Pullar, B.J., Kirkness, E.F., Kruglyak, L., & Ostrander, E.A. (2004). Extensive and breed-specific linkage disequilibrium in Canis familiaris. Genome research, 14(12), 2388–2396.

Szklarczyk, D., Kirsch, R., Koutrouli, M., Nastou, K., Mehryary, F., Hachilif, R., Gable, A.L., Fang, T., Doncheva, N.T., & Pyysalo, S. (2023). The STRING database in 2023: protein–protein association networks and functional enrichment analyses for any sequenced genome of interest. Nucleic acids research, 51(D1), D638–D646.

Tye, K.M., Mirzabekov, J.J., Warden, M.R., Ferenczi, E.A., Tsai, H.-C., Finkelstein, J., Kim, S.-Y., Adhikari, A., Thompson, K.R., & Andalman, A.S. (2013). Dopamine neurons modulate neural encoding and expression of depression-related behaviour. Nature, 493(7433), 537–541.

Tye, K.M., Prakash, R., Kim, S.-Y., Fenno, L.E., Grosenick, L., Zarabi, H., Thompson, K.R., Gradinaru, V., Ramakrishnan, C., & Deisseroth, K. (2011). Amygdala circuitry mediating reversible and bidirectional control of anxiety. Nature, 471(7338), 358–362.

Van Alstyne, M., Tattoli, I., Delestrée, N., Recinos, Y., Workman, E., Shihabuddin, L.S., Zhang, C., Mentis, G.Z., & Pellizzoni, L. (2021). Gain of toxic function by long-term AAV9-mediated SMN overexpression in the sensorimotor circuit. Nature Neuroscience, 24(7), 930–940. 10.1038/s41593-021-00827-3

Van de Sande, B., Flerin, C., Davie, K., De Waegeneer, M., Hulselmans, G., Aibar, S., Seurinck, R., Saelens, W., Cannoodt, R., Rouchon, Q., et al. (2020). A scalable SCENIC workflow for single-cell gene regulatory network analysis. Nature Protocols, 15(7), 2247–2276. 10.1038/s41596-020-0336-2

Walf, A.A., & Frye, C.A. (2007). The use of the elevated plus maze as an assay of anxiety-related behavior in rodents. Nature protocols, 2(2), 322–328.

Wallis, N.J., McClellan, A., Mörseburg, A., Kentistou, K.A., Jamaluddin, A., Dowsett, G.K., Schofield, E., Morros-Nuevo, A., Saeed, S., & Lam, B.Y. (2025). Canine genome-wide association study identifies DENND1B as an obesity gene in dogs and humans. Science, 387(6741), eads2145.

Wang, C., Wallerman, O., Arendt, M.-L., Sundström, E., Karlsson, Å., Nordin, J., Mäkeläinen, S., Pielberg, G.R., Hanson, J., & Ohlsson, Å. (2021). A novel canine reference genome resolves genomic architecture and uncovers transcript complexity. Communications Biology, 4(1), 185.

Wang, D., Liu, S., Warrell, J., Won, H., Shi, X., Navarro, F.C.P., Clarke, D., Gu, M., Emani, P., Yang, Y.T., et al. (2018). Comprehensive functional genomic resource and integrative model for the human brain. Science, 362(6420), eaat8464. 10.1126/science.aat8464

Wolf, F.A., Angerer, P., & Theis, F.J. (2018). SCANPY: large-scale single-cell gene expression data analysis. Genome biology, 19(1), 15.

Xu, Z., Zhu, J., Ma, Z., Zhen, D., & Gao, Z. (2025). Combined Bulk and Single-Cell Transcriptomic Analysis to Reveal the Potential Influences of Intestinal Inflammatory Disease on Multiple Sclerosis. Inflammation, 48(4), 2367–2386. 10.1007/s10753-024-02195-z

Yang, L., Guo, C., Zheng, Z., Dong, Y., Xie, Q., Lv, Z., Li, M., Lu, Y., Guo, X., & Deng, R. (2025). Stress dynamically modulates neuronal autophagy to gate depression onset. Nature, 641(8062), 427–437.

Yao, Z., Van Velthoven, C.T., Kunst, M., Zhang, M., McMillen, D., Lee, C., Jung, W., Goldy, J., Abdelhak, A., & Aitken, M. (2023). A high-resolution transcriptomic and spatial atlas of cell types in the whole mouse brain. Nature, 624(7991), 317–332.

Yu, Y., Wu, Y., Zhao, H., Zhou, Z., Bai, J., Zhang, Y., Zhang, F., Zhou, B., Wang, J., & Zhang, Y.-P. (2026). Genetic basis of canine separation-related behavior: putative enhancer-dependent regulation of SLC32A1 and conservation in human psychiatric disorders. Journal of Genetics and Genomics. 10.1016/j.jgg.2026.03.003

Zapata, I., Hecht, E.E., Serpell, J.A., & Alvarez, C.E. (2020). Genome scans of dog behavior implicate a gene network underlying psychopathology in mammals, including humans. BioRxiv, 2020.2007. 2019.211078.

Zhang, M., Li, X., Herman, J.G., Gao, A., Wang, Q., Yao, Y., Shen, F., He, K., & Guo, M. (2024). Methylation of NRIP3 Is a synthetic lethal marker for combined PI3K and ATR/ATM inhibitors in colorectal cancer. Clinical and Translational Gastroenterology, 15(3), e00682.

Zhang, Q., Liu, H., & Bu, F. (2021). High performance of a GPU-accelerated variant calling tool in genome data analysis. BioRxiv, 2021.2012.2012.472266. 10.1101/2021.12.12.472266

Zhang, W., Lu, Y., Shen, R., Wu, Y., Liu, C., Fang, X., Zhang, L., Liu, B., & Rong, L. (2025). Inhibiting ceramide synthase 5 expression in microglia decreases neuroinflammation after spinal cord injury. Neural Regeneration Research, 20(10), 2955–2968.

Zhou, X., & Stephens, M. (2012). Genome-wide efficient mixed-model analysis for association studies. Nature Genetics, 44(7), 821–824.

Ziemann, A.E., Allen, J.E., Dahdaleh, N.S., Drebot, I.I., Coryell, M.W., Wunsch, A.M., Lynch, C.M., Faraci, F.M., Howard, M.A., & Welsh, M.J. (2009). The amygdala is a chemosensor that detects carbon dioxide and acidosis to elicit fear behavior. Cell, 139(5), 1012–1021.

