## Supplementary Figures for "GWAS and multi-omics uncover the *SLC6A3* locus for canine stranger-directed social anxiety and implicate *NRIP3* in dopaminergic network regulation"

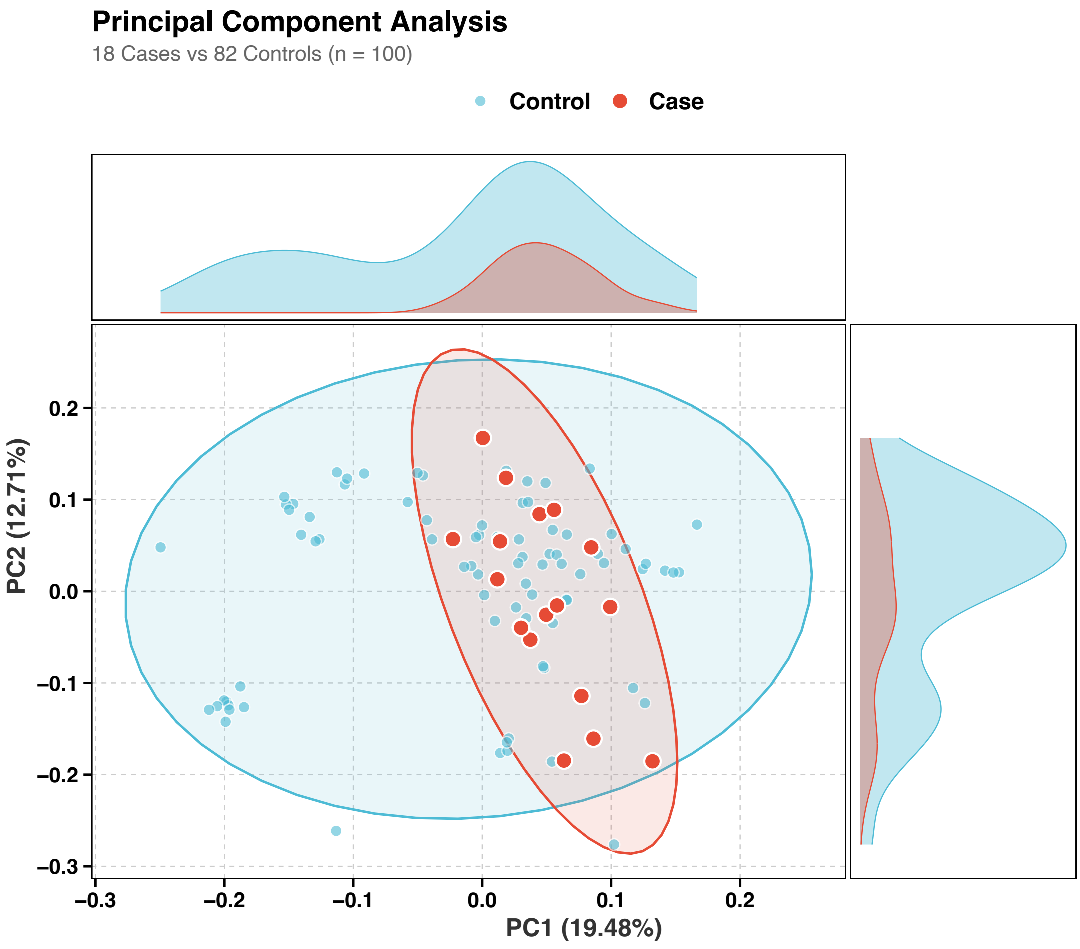


**Figure S1.** Principal component analysis of genetic population structure in Chinese Kunming dogs. PCA was performed using 174,290 LD-pruned variants from the final GWAS cohort of 100 Chinese Kunming dogs, including 18 dogs with social anxiety-like behavior and 82 behaviorally normal controls. PC1 and PC2 explained 19.5% and 12.7% of the genetic variance, respectively. Cases and controls were broadly intermingled in the PCA space, indicating no obvious separation by case-control status. PC1 and PC2 were included as covariates in the GEMMA linear mixed model to control for population structure.

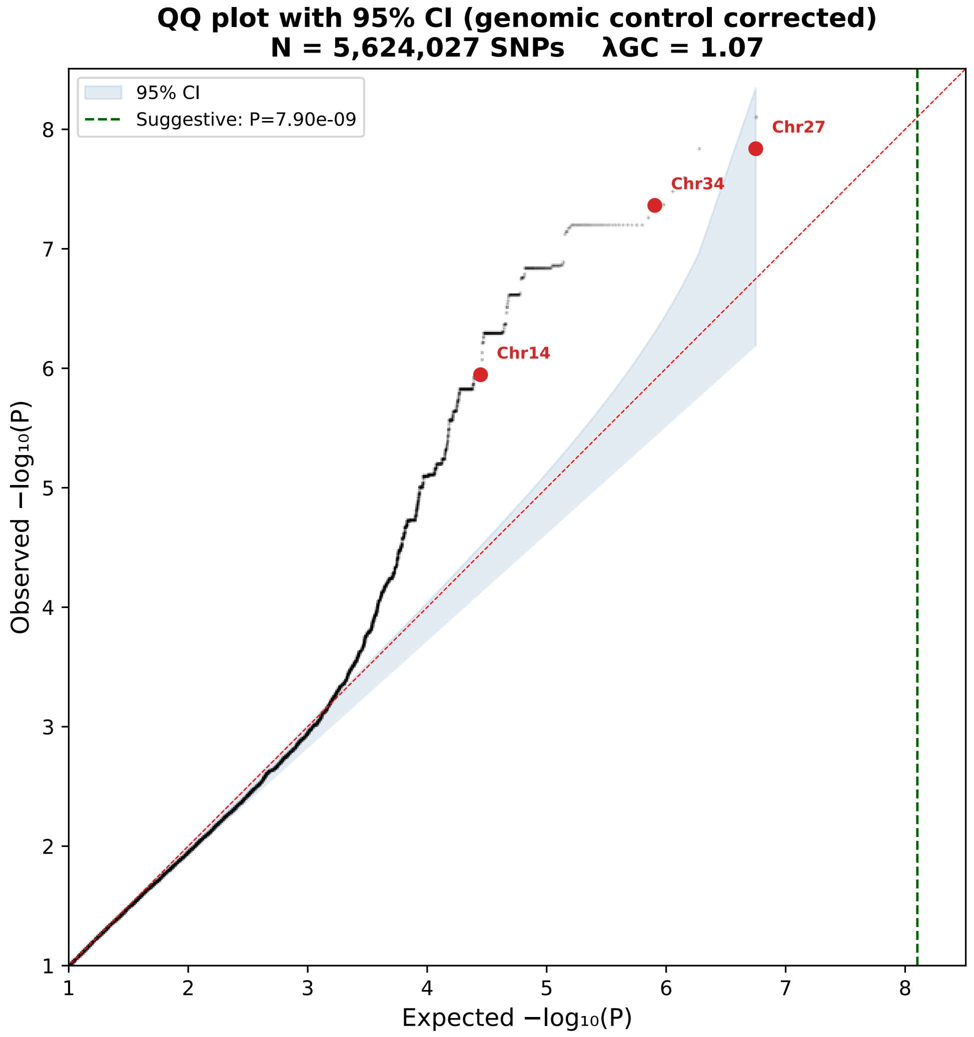


**Figure S2.** Quantile-quantile plot of the canine GWAS for social anxiety-like behavior. The QQ plot compares observed and expected P values from the GEMMA linear mixed model GWAS in 100 Chinese Kunming dogs. The genomic inflation factor was λGC = 1.07, indicating limited inflation and adequate control of population stratification and cryptic relatedness after inclusion of PC1, PC2, sex, and the genomic relationship matrix in the association model. The deviation at the tail reflects the strongest association signals rather than systematic genome-wide inflation.

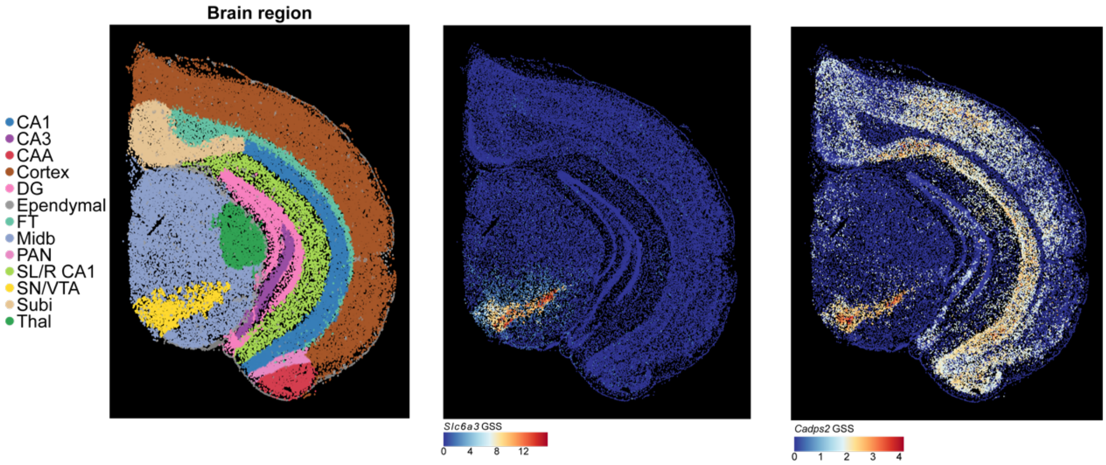


**Figure S3. gsMap-enhanced spatial transcriptomic analysis of Slc6a3 and Cadps3 in the mouse brain.** Left, anatomical annotation of major brain regions. Middle and right, spatial distributions of gsMap-derived GSS for *Slc6a3* and *Cadps3*, respectively. Both genes show prominent enrichment in the SN/VTA region, with warmer colors indicating higher GSS values.

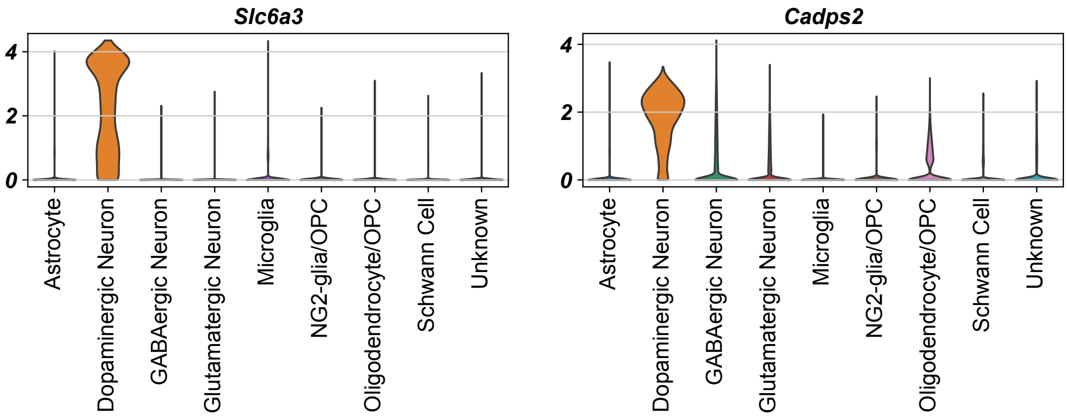


**Figure S4. Cell-type-specific expression distributions of Slc6a3 and Cadps2 in the mouse midbrain single-cell dataset.** Violin plots show the expression levels of *Slc6a3* and *Cadps2* across annotated cell populations, including astrocytes, dopaminergic neurons, GABAergic neurons, glutamatergic neurons, microglia, NG2-glia/OPCs, oligodendrocytes/OPCs, Schwann cells, and unknown cells. *Slc6a3* showed highly selective expression in dopaminergic neurons, consistent with its role as a canonical dopaminergic marker. *Cadps2* was also enriched in dopaminergic neurons, although lower expression was observed in several non-dopaminergic cell populations.

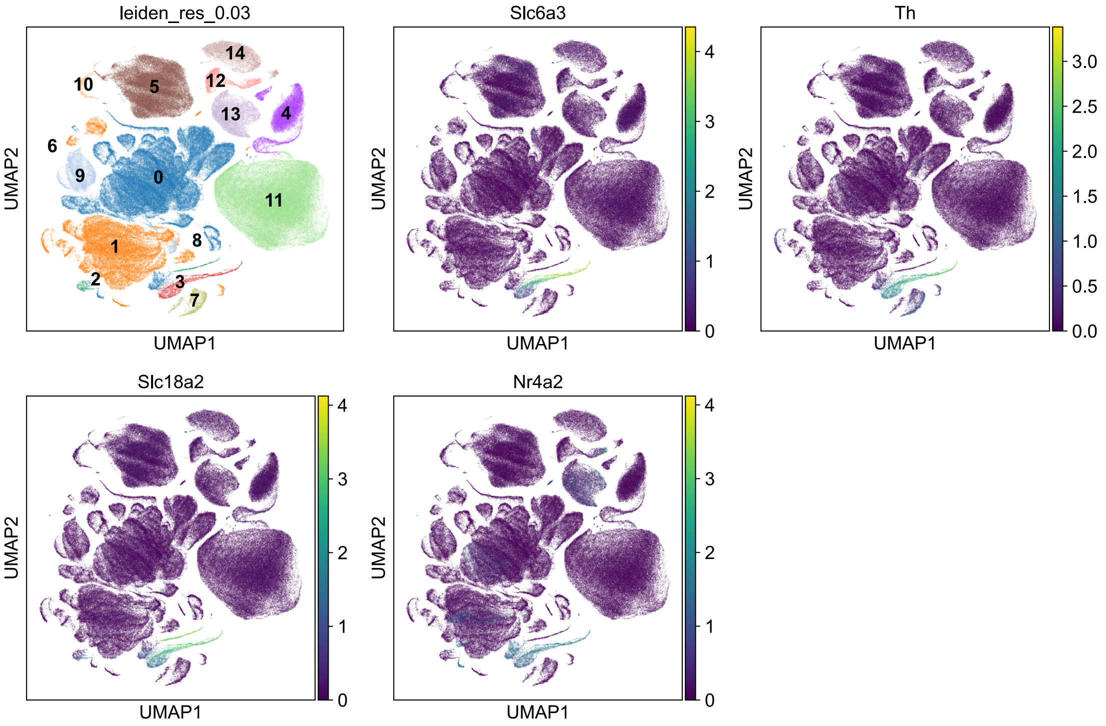


**Figure S5. Expression of canonical dopaminergic neuron markers used for cell annotation.** UMAP plots showing the expression of canonical dopaminergic neuron markers (*Slc6a3*, *Th*, *Slc18a2*, and *Nr4a2*) used for identifying dopaminergic neurons. The consistent expression of these markers supports the annotation of this neuronal population.

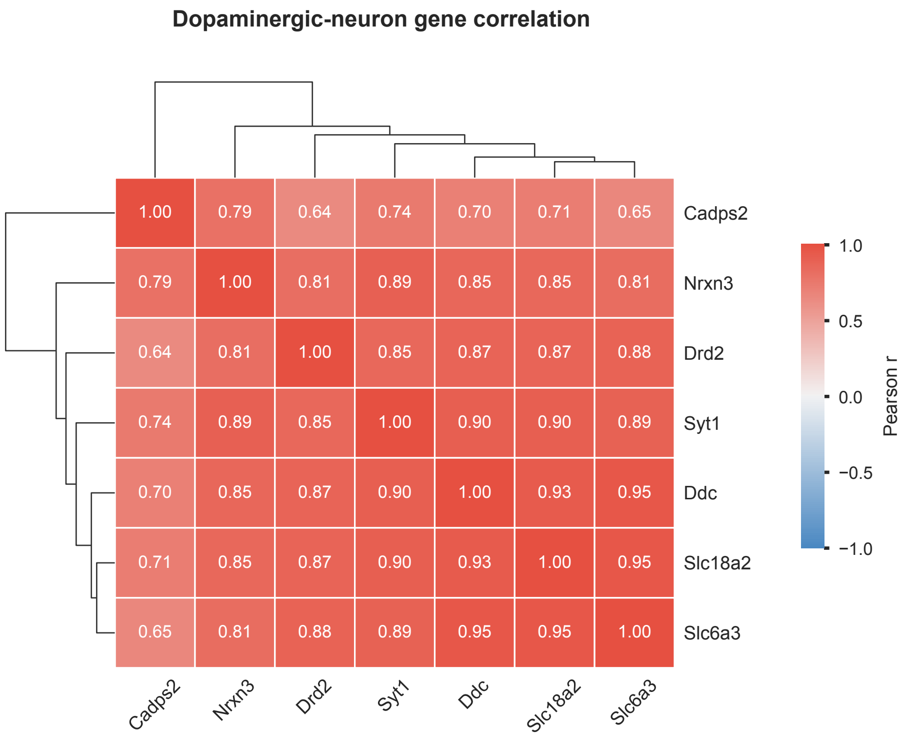


**Figure S6. Co-expression structure of dopamine-related genes in dopaminergic neurons.** Clustered heatmap showing Pearson correlation coefficients among *Slc18a2*, *Ddc*, *Slc6a3*, *Cadps2*, *Drd2*, *Syt1* and *Nrxn3* based on raw single-cell expression counts from annotated dopaminergic neurons. Rows and columns were ordered by hierarchical clustering using Pearson distance (1 - r). Values inside the heatmap indicate Pearson's r. The analysis included 3,353 dopaminergic neurons.

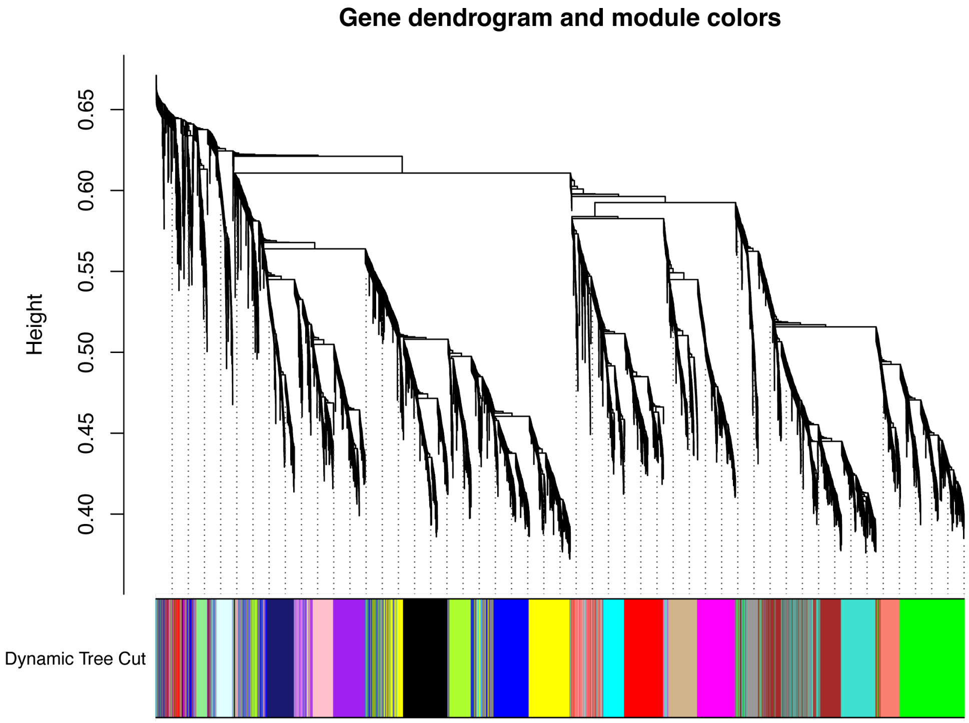


**Figure S7. WGCNA reveals conserved co-expression of *SLC6A3* and *CADPS2* in the human VTA.** Both genes are robustly assigned to the same co-expression network (yellow module) based on public transcriptomic data from 24 human VTA samples.

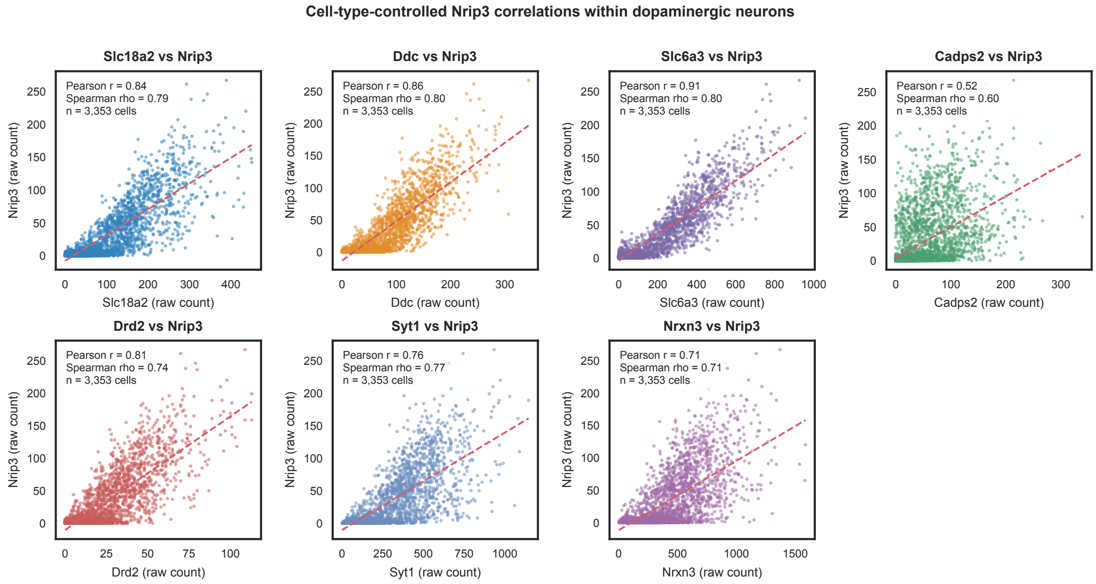


**Figure S8. Cell-type-controlled correlation between *Nrip3* and dopamine-related genes.** Scatter plots showing pairwise correlations between *Nrip3* and *Slc18a2*, *Ddc*, *Slc6a3*, *Cadps2*, *Drd2*, *Syt1* or *Nrxn3* in annotated dopaminergic neurons. Each point represents one cell, and expression values are shown as raw counts. Dashed lines indicate least-squares linear regression fits. Pearson and Spearman correlation coefficients are shown in each panel; P values were not calculated. The analysis included 3,353 dopaminergic neurons.

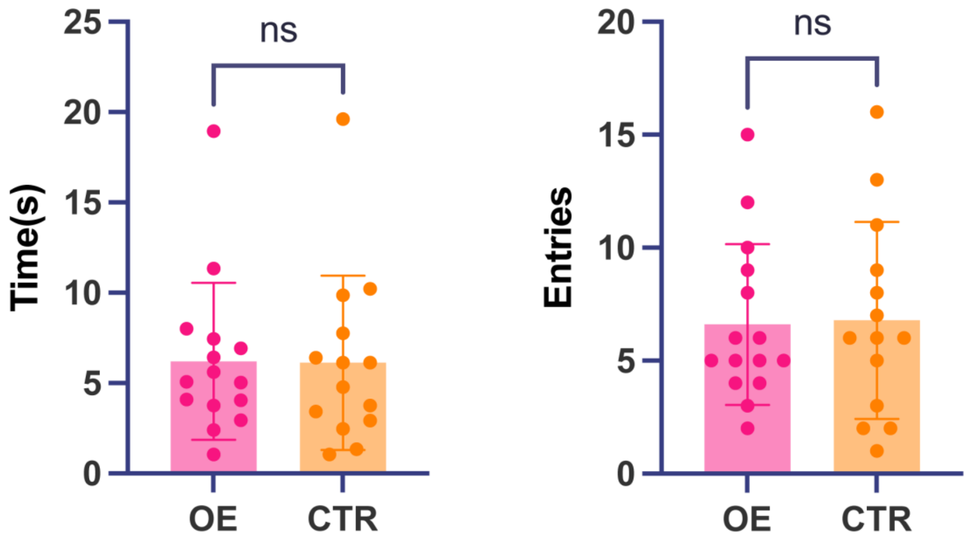


**Figure S9. Overexpression does not alter anxiety-like behavior in the Open Field Test.** The time spent in the center zone and the number of entries into it were comparable between the overexpression and control mice.

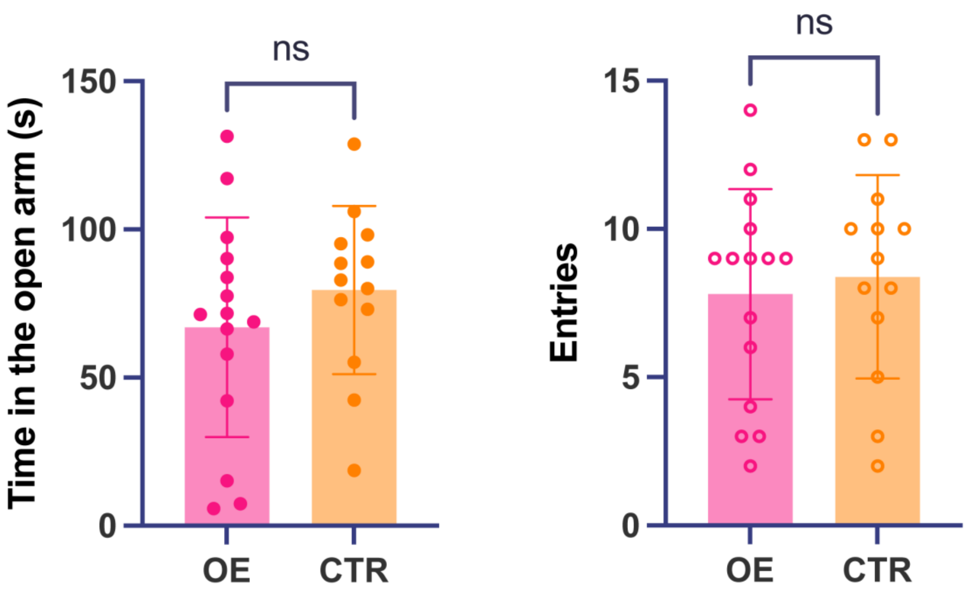


**Figure S10. Elevated Plus Maze (EPM) assessment of anxiety-related behavior.** No significant differences were observed between the two groups in the total number of entries or time spent in the open arms.

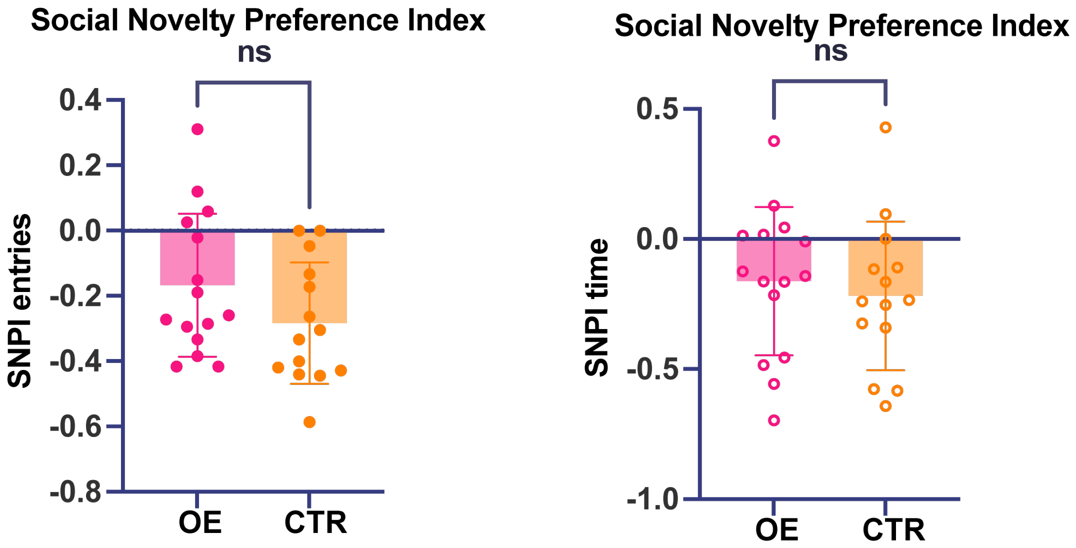


**Figure S11. Social novelty preference in the three-chamber social interaction test.** Social novelty preference time index and social novelty preference entry index during phase 3 of the test. *Nrip3*-overexpression mice did not differ significantly from controls in either time preference (OE, n = 15; CTR, n = 14; Welch's t = 0.5303, df = 26.86, P = 0.6003) or entry preference (Welch's t = 1.547, df = 26.78, P = 0.1335).

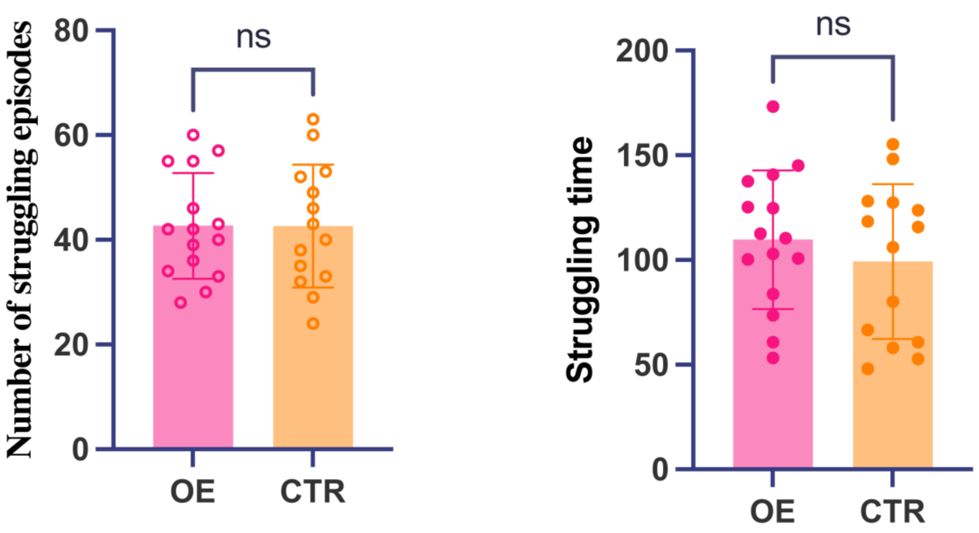


**Figure S12. Tail Suspension Test (TST) assessment of depressive-like behavior.** No significant differences were observed between the two groups in the total struggling time or the number of struggles.
